# Three hallmarks of malaria-induced selection in human genomes

**DOI:** 10.1101/2020.07.07.170092

**Authors:** Jacob A. Tennessen, Manoj T. Duraisingh

**Affiliations:** Harvard T.H. Chan School of Public Health, Boston, MA USA

## Abstract

Malaria has plausibly been the single strongest selective pressure on our species. Many of the best-characterized cases of adaptive evolution in humans are in genes tied to malaria resistance. However, the complex evolutionary patterns at these genes are poorly captured by standard scans for non-neutral evolution. Here we present three new statistical tests for selection based on population genetic patterns that are observed more than once among key malaria resistance loci. We assess these tests using forward-time evolutionary simulations and apply them to global whole-genome sequencing data from humans, and thus we show that they are effective at distinguishing selection from neutrality. Each test captures a distinct evolutionary pattern, here called Divergent Haplotypes, Repeated Shifts, and Arrested Sweeps, associated with a particular period of human prehistory. We clarify the selective signatures at known malaria-relevant genes and identify additional genes showing similar adaptive evolutionary patterns. Among our top outliers, we see a particular enrichment for genes involved in erythropoiesis and for genes previously associated with malaria resistance, consistent with a major role for malaria in shaping these patterns of genetic diversity. Polymorphisms at these genes are likely to impact resistance to malaria infection and contribute to ongoing host-parasite coevolutionary dynamics.

## Introduction

Malaria, a major global infectious disease caused by *Plasmodium* parasites and spread by mosquitoes, has been one of the most important selective pressures on the human lineage (Ebel et al. 2017). Bolstered by the intimate coevolutionary history between humans and *Plasmodium* and the severe pathology of malaria, several of the strongest signatures of selection in the human genome center on genes that impact malaria resistance. These genes include *ABO* (A/B/O blood group; Ségurel et al. 2012, 2013), the cluster of *GYPA, GYPB*, and *GYPE* (here abbreviated to *GYPA/B/E*, glycophorin Dantu blood group; Leffler et al. 2017), *ACKR1* (Duffy antigen; Hamblin et al. 2002; King et al. 2011; Chittoria et al. 2012), *CR1* (Knops blood group; Tham et al. 2010; Prajapati et al. 2019), *HBB* (hemoglobin B; Allison 1954; Laval et al. 2019), and *G6PD* (glucose-6-phosphate dehydrogenase; Ruwende et al. 1995; Tishkoff et al. 2001). Malaria remains a major selection pressure to this day, with 200 million cases annually, leading to over 400,000 deaths (Miller et al. 2002; WHO 2019).

The adaptive signatures wrought by *Plasmodium* on humans are useful to characterize and study, for two reasons. First, evolutionary signatures have been critical for finding new malaria-relevant genes (*HBB*, Allison 1954; *GYPA/B/E*, Malaria Genomic Epidemiology Network 2015). There are likely more large-effect genes to be found, as over half of the heritability in malaria resistance (h^2^ ~24%) remains unexplained (Mackinnon et al. 2005; Malaria Genomic Epidemiology Network 2019) and there is substantial geographic heterogeneity in the genetic basis of resistance (Leffler et al. 2017). Progress toward malaria elimination has stalled in recent years, prompting the need for new treatments (White et al. 2014; WHO 2019). Understanding the genetic basis of malaria resistance can pave the way for therapeutics that target host-parasite molecular interactions (Cowman et al. 2017) and inform precision medicine. Second, malaria resistance genes present a robust model system to develop and assess statistical tests for selection, given their striking evolutionary signatures and well-documented phenotypic effects (Malaria Genomic Epidemiology Network 2019). Such tests may be broadly applicable to study other selective pressures, for example in other host-parasite systems.

Genome-wide scans for selection in humans are now routine (Fan et al. 2016), but they have poor replicability such that different methods produce very different lists of candidate genes. This is true for positive selection (Akey 2009) and may be worse for balancing selection: among six recent studies that scan the genome for balancing selection in Africans or African Americans, the proportion of identified candidate selection targets that are shared between any two scans ranges from 0 to 9% (Andrés et al. 2009; Leffler et al. 2013; DeGiorgio et al. 2014; Siewert and Voight 2017; Bitarello et al. 2018; Cheng and DeGiorgio 2019). For some genes, selection has been effectively validated phenotypically because allelic effects on infection or fitness have been demonstrated. However, while these known causal genes (e.g. *HBB*, *ABO*, *G6PD*) do show unusual and presumably non-neutral population genetic patterns, they overlap poorly with genome-wide selection scans, suggesting that existing tests are underpowered to detect true positives. There is both a need for analytical tools that can better distinguish biologically meaningful polymorphism from neutral polymorphism, and an opportunity to leverage these functionally validated loci to guide the development of such tools.

In this paper, we focus on three population genetic patterns that are common in malaria-relevant genes but are poorly approximated by existing statistical tests for selection (Table 1; Figure 1). We call these patterns Divergent Haplotypes, Repeated Shifts, and Arrested Sweeps. Each is represented by two exemplar loci (Table 1). We develop new ways to summarize population genetic data that readily distinguish these signatures from the neutral background, and assess these statistics using simulations. In order to diversify the character set for population genetics beyond the heavily used Latin and Greek alphabets, each statistic is denoted by an emoji, as these Unicode characters are now a standard component of publishing software and have been underutilized in science (O’Reilly-Shah et al. 2018): Dango 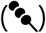, Trident 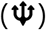, and Pause 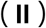. Finally, we apply our tests to population genomic data (1000 Genomes Project Consortium 2015) to evaluate their ability to detect known malaria-relevant genes and to identify new candidate genes.

**Table 1.**
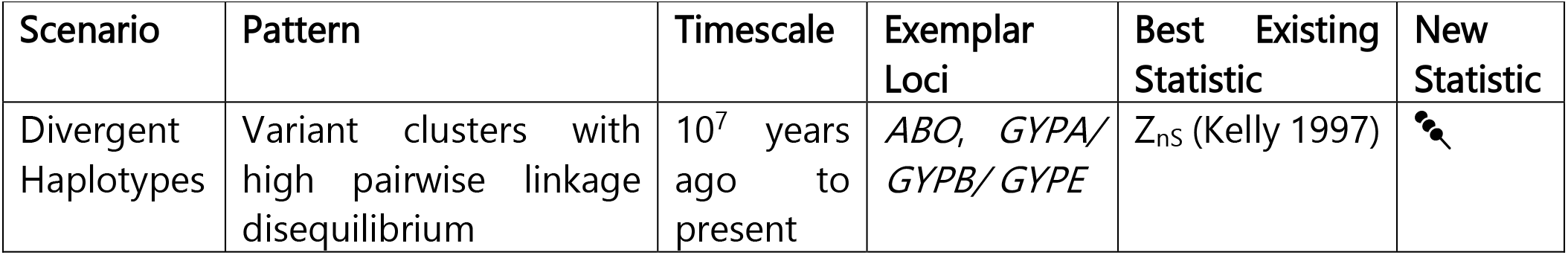
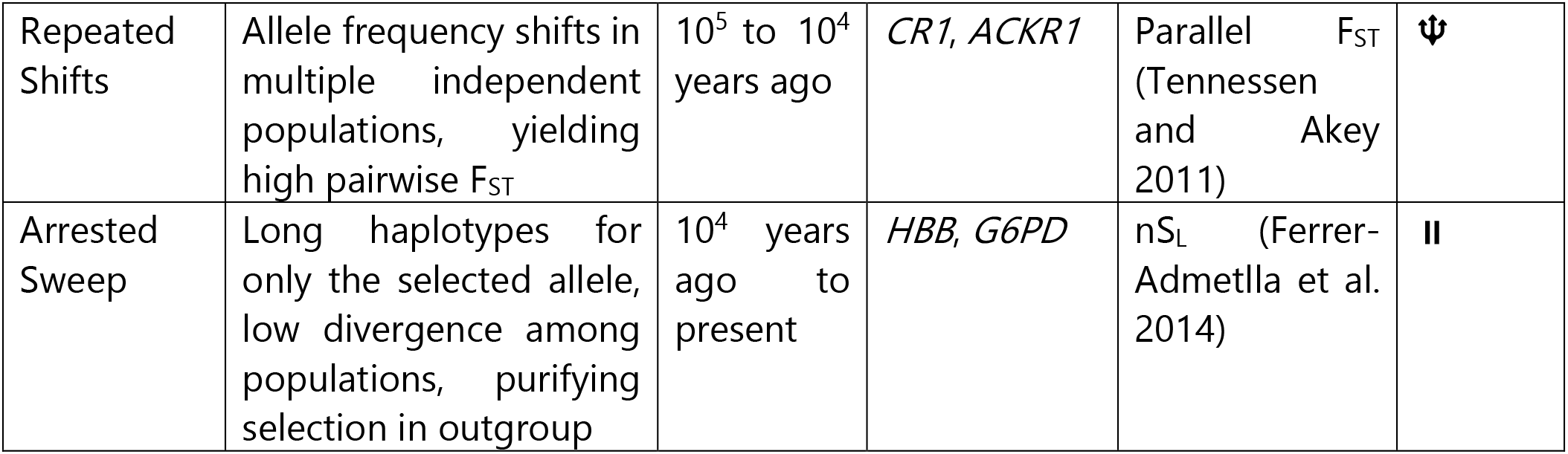
Framework for detecting three adaptive evolutionary scenarios.

**Figure 1.**
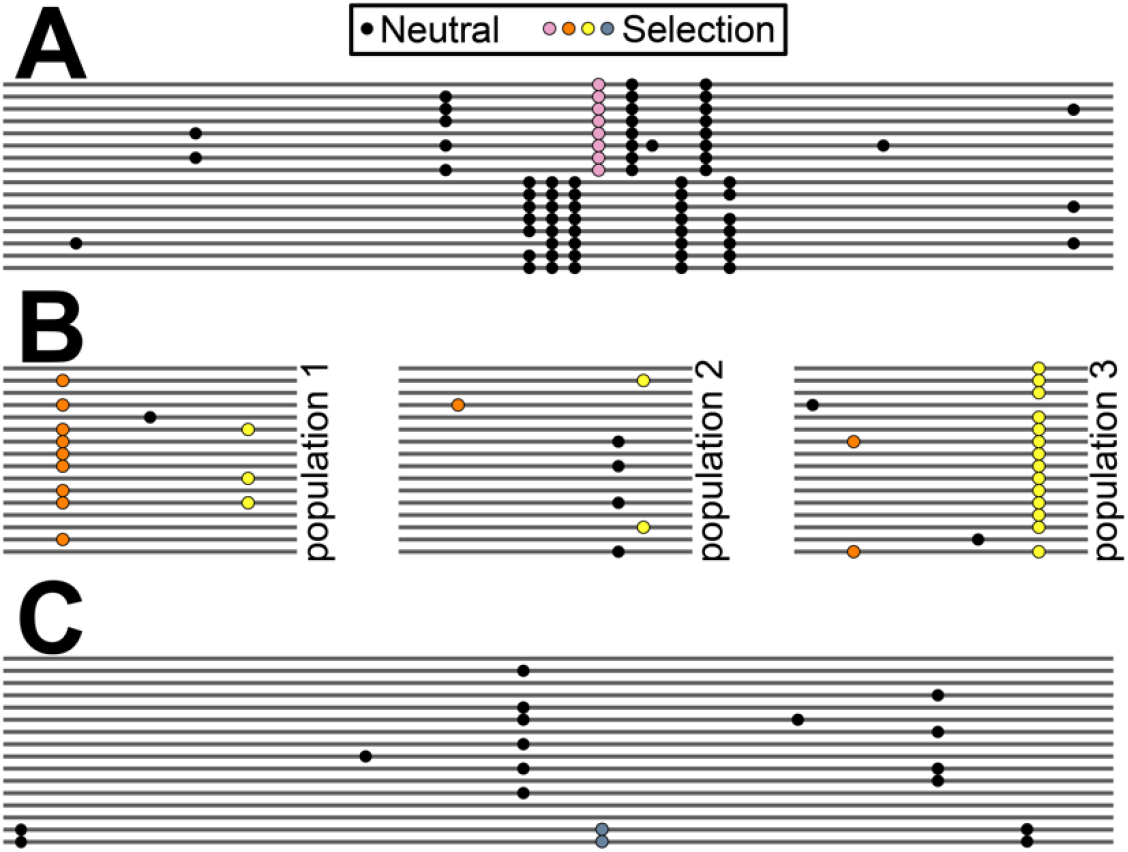
Three complex adaptive scenarios observed repeatedly among malaria-relevant genes. Lines represent chromosomes. Circles are derived alleles at polymorphic sites (black = neutral; colored = adaptive). (A) Divergent Haplotypes. A dense cluster of variants in high linkage disequilibrium occurs within a narrow genomic window surrounding a balanced polymorphism. (B) Repeated Shifts. All three pairwise population comparisons show unusually high divergence at one or more variants, suggesting repeated bouts of positive selection. (C) Arrested Sweep. A beneficial mutation swept up a long haplotype but stopped at relatively low frequency.

## New Approaches

### Divergent Haplotypes and 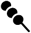

Divergent Haplotypes are observed at *ABO* and *GYPA/B/E*. At these loci, distinct haplotypes have been maintained by balancing selection for millions of years, as evidenced by non-human primates sharing the polymorphism (Ségurel et al. 2012; Leffler et al. 2013; Malaria Genomic Epidemiology Network 2015). While trans-species polymorphisms represent strong evidence for selection, they likely constitute a small minority of balanced polymorphisms as they require consistent selection for very long periods across distinct ecological niches. They are therefore unreliable as a comprehensive metric for selection targets. *ABO* and *GYPA/B/E* do show non-neutral patterns within human populations, but the signal is relatively weak for commonly used tests (Stajich and Hahn 2005; Leffler et al. 2017). As a result, scans for balancing selection fail to identify either of these genes as outliers (e.g. Andrés et al. 2009; DeGiorgio et al. 2014; Bitarello et al. 2018), unless they incorporate non-human polymorphism (Leffler et al. 2013; Chang and DeGiorgio 2019). However, intraspecies haplotype structure alone may convey a signal of selection if tested for directly (Figure 1A).

We developed and evaluated a new test for Divergent Haplotypes. Old, balanced haplotypes accumulate mutations which are protected from genetic drift, leading to increased sequence divergence between the haplotypes (Figure 1A). This pattern is disrupted by recombination, so it can typically be observed only across small genetic distances. Clusters of closely adjacent variants in high linkage disequilibrium (LD) and similar minor allele frequencies (MAF) are thus a signature of balancing selection (Siewert and Voight 2017), as envisioned by Kelly (1997) with the Z_nS_ statistic. However, Z_nS_ is highly sensitive to individual rare variants, which will typically not show high LD with any other variant. Rare variants may or may not be observed depending on stochasticity and sample size, making Z_nS_ a very noisy statistic. We therefore define a new test statistic which sums across LD correlations rather than averaging them: 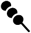 (equation 3). For a target variant, 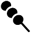 is the sum of LD correlations with all other variants within a distance of *g* bp. Using 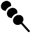, rare variants have a negligible effect, and the statistic is maximized if there are a large number of variants within a narrow region in high LD with each other. There is no upper limit to 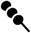, and its typical range for a given population will depend on overall levels of nucleotide diversity and LD. Therefore, unusual values of 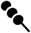 are defined in comparison to the genome-wide average.

### Repeated Shifts and 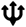

Repeated Shifts are a form of parallel adaptive divergence, which occurs when positive selection repeatedly causes rapid allele frequency change, resulting in high F_ST_ (defined here following Weir and Cockerham 1984), at the same locus in geographically distinct populations (Figure 1B). High divergence between independent pairs of populations is unlikely to occur more than once unless driven by natural selection (Tennessen and Akey 2011). Notably, some of the strongest human instances of parallel adaptive divergence occur in genes with large effects on malaria resistance. *ACKR1* is not only the single most divergent gene between Africa and Europe, but it is also among the most divergent genes between Europe and Asia. This is due to near-fixation of the Duffy-null allele in sub-Saharan Africa, and independent selection for the Fy^a^ allele in Asia (Hamblin et al. 2002; King et al. 2011; Chittoria et al. 2012). Similarly, *CR1* is also divergent in both Africa-Europe and Asia-Europe comparisons due to positive selection (Prajapati et al. 2019). Our previous genome-wide scan for parallel adaptive divergence (Tennessen and Akey 2011) sought repeated, phylogenetically independent shifts occurring at the same single-nucleotide variant among a set of four populations. Such strict criteria miss *ACKR1* and *CR1*. These genes show changes at different variants in different population comparisons, and these comparisons are nested rather than truly independent (i.e. high Africa v. Eurasia divergence at some variants, and high Africa/Europe v. Asia divergence at other variants). A revised approach might have enhanced power to detect such cases.

We developed and evaluated a new test for Repeated Shifts. Our goal was to detect narrow genomic windows showing unusually high F_ST_ in all three pairwise comparisons among three populations, at the same or different variants (Figure 1B). Such population triplets are not phylogenetically independent as in Tennessen and Akey (2011), but they cannot be explained without multiple bouts of positive selection. Our approach is based on inversely ranking genomic windows of size *g* bp based on F_ST_ and finding windows in which all three pairwise ranks are relatively extreme. Our test statistic 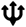 (equation 4) is a squared average between the lowest (*L*) and highest (*H*) F_ST_ ranks scaled by the number of windows examined, which approximates a p-value reflecting the probability of observing the data if there is no parallel selection acting. However, even values of 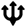 that are not individually significant may provide evidence for selection if they are among the most extreme values in the genome.

### Arrested Sweeps and 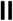

The Arrested Sweep pattern is observed at *HBB* and *G6PD*. At both of these loci, an allele protective against malaria arose in Africa 5-25 kya (Tishkoff et al. 2001; Shriner and Rotimi 2018; Laval et al. 2019) and rapidly increased in frequency. At both loci, the protective allele then stopped spreading and has been maintained at about 10% frequency because it conveys a physiological disadvantage when homozygous/hemizygous (Figure 1C). Because of the low frequency of the derived allele, these genes are missed by selection scans that seek intermediate frequency alleles or high intercontinental F_ST_. Because the recent timescale has precluded the accumulation of elevated nucleotide diversity or trans-species polymorphisms, these genes are missed by scans for ancient haplotypes. Thus, despite being canonical examples of adaptive polymorphism, these genes are almost never detected in genome-wide scans for partial sweeps or balancing selection (e.g. Voight et al. 2006; Akey 2009; Andrés et al. 2009; Leffler et al. 2013; DeGiorgio et al. 2014; Siewert and Voight 2017; Bitarello et al. 2018; Cheng and DeGiorgio 2019).

We developed and evaluated a new test for Arrested Sweeps. This test seeks recently arisen variants which are beneficial when heterozygous but strongly deleterious otherwise, as with *HBB* and *G6PD* (Figure 1C). This test is conducted on a single target population. It also requires several other populations hypothesized to experience similar selective pressures; along with the target population, these constitute the “ingroup”. Finally, it requires an “outgroup” population in which there is no heterozygote advantage to the derived variant, only the deleterious effect (as with any population where malaria does not occur). There are two evolutionary components of an Arrested Sweep: the sweep (positive selection) and the arrest (balancing and purifying selection).

The first evolutionary step is a partial positive selective sweep. A sweep leaves a signal of low polymorphism and long-range LD within the beneficial haplotypic lineage, as captured by extended haplotype homozygosity statistics like iHS (Voight et al. 2006). However, iHS does not readily detect *HBB* or *G6PD* (Voight et al. 2006), and related statistics target other evolutionary scenarios like soft sweeps or sweeps near completion (Garud et al. 2015; Sabeti et al. 2007). The statistic nS_L_ does implicate *HBB* (Ferrer-Admetlla et al. 2014; Laval et al. 2019), but this statistic, like any based on haplotype homozygosity, is sensitive to relatedness among samples and new mutations or sequencing errors that disrupt the otherwise perfect similarity among haplotypes. However, the signal of a sweep may extend beyond the range of haplotype homozygosity, in the form of reduced, but not necessarily nonzero, nucleotide diversity linked to the swept haplotype. In other words, individuals homozygous for a swept allele have fewer total heterozygous sites than individuals homozygous for the ancestral allele.

The second evolutionary step, which distinguishes an Arrested Sweep from an ongoing partial sweep, is balancing selection maintaining the polymorphism at the optimal frequency in the ingroup while purifying selection excludes it from the outgroup. Assessing this signal starts with F_ST_ within the ingroup. An arrested sweep maintained at constant frequency by balancing selection will show low F_ST_ among populations that are experiencing the same selection pressure. An ongoing sweep should not show this pattern and may even show unusually high F_ST_ if the sweep has proceeded farther in some populations than others. In addition, purifying selection acts in the outgroup where there is no heterozygote advantage, so the MAF in the outgroup should be very close to zero.

Our final test statistic 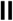 (equation 7) is a product of metrics that capture these steps: positive sweep in the target population, balancing selection across the ingroup, and purifying selection in the outgroup. There is no upper limit to 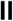 and unusual values of 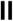 are defined in comparison to the genome-wide average.

## Results

### 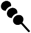 in simulations

Simulation results show that 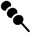 values are typically higher for balanced polymorphisms, relative to neutral polymorphisms, under a wide range of parameters (Supp Figure 1). Intermediate *g* values (500 to 1000 bp) were optimal to minimize the overlap between selection and neutral windows. The power of 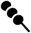 is maximized when recombination rates are low, the polymorphism is old, and mutation rates surrounding balanced polymorphisms are similar to those in neutral regions. Skewing the expected MAF had little effect, in contrast to many common tests for balancing selection that seek intermediate-frequency variants. For *g* of 500 and human-relevant parameters, variants with approximately 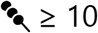 should be enriched for true balanced polymorphisms, though with inevitable false positives and false negatives.

### 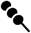 in human population data

In our focal population YRI (Yoruba in Ibadan, Nigeria), 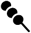 with *g* of 500 ranged from 0 to 59.8 (median = 0.2, 95% interval = 0.0 to 5.9; Figure 2A). Only 0.56% of variants had 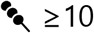. Variants in or near the HLA accounted for 15% of variants with 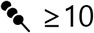, a majority (66%) of variants in the top 0.05% 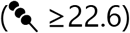 and the 58 highest variants (Figure 2B). There are 243 coding genes with at least one exonic 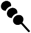 value ≥10, including *ABO* (Figure 2C). The glycophorins do not show a strong exonic signal but many high-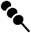 variants, including two reaching the top 0.05% threshold, occur in the intergenic region between *GYPE* and *FREM3* where the strongest signal of selection and disease-association of the glycophorin cluster has previously been detected (Leffler et al. 2013; Malaria Genomic Epidemiology Network 2015; Figure 2D). These results suggest that 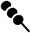 does capture the intended empirical selection signal. The top 50 genes outside of the HLA region (Table 2; Supp Table 1; Figure 2 E,F), based on 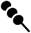 in exons or within 1 kb upstream, all showed 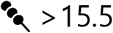 (top 0.15% of variants).

**Table 2.**
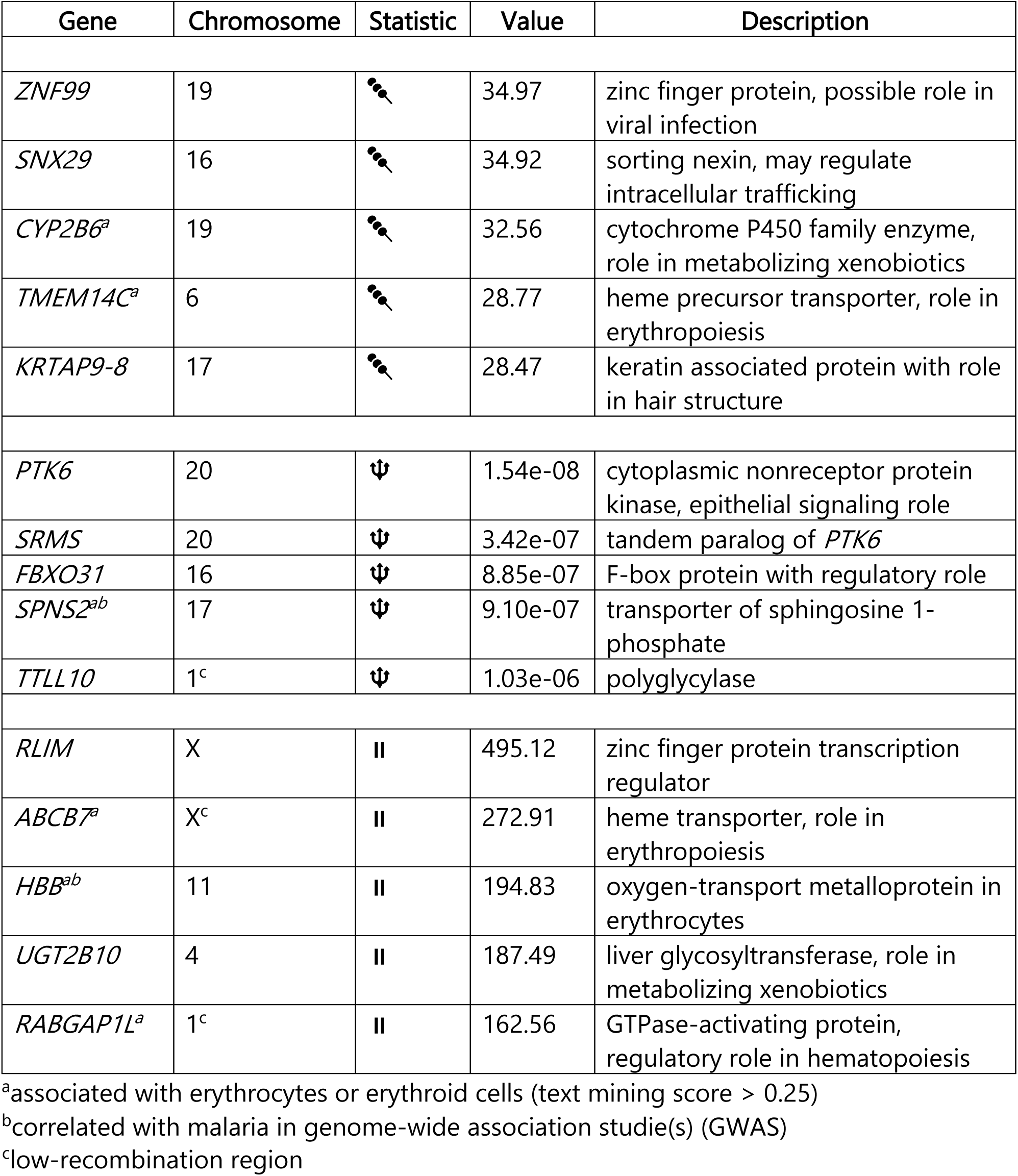
Top five genes for each statistic when occurring in exons or within 1 kb upstream, excluding the HLA region.

**Figure 2.**
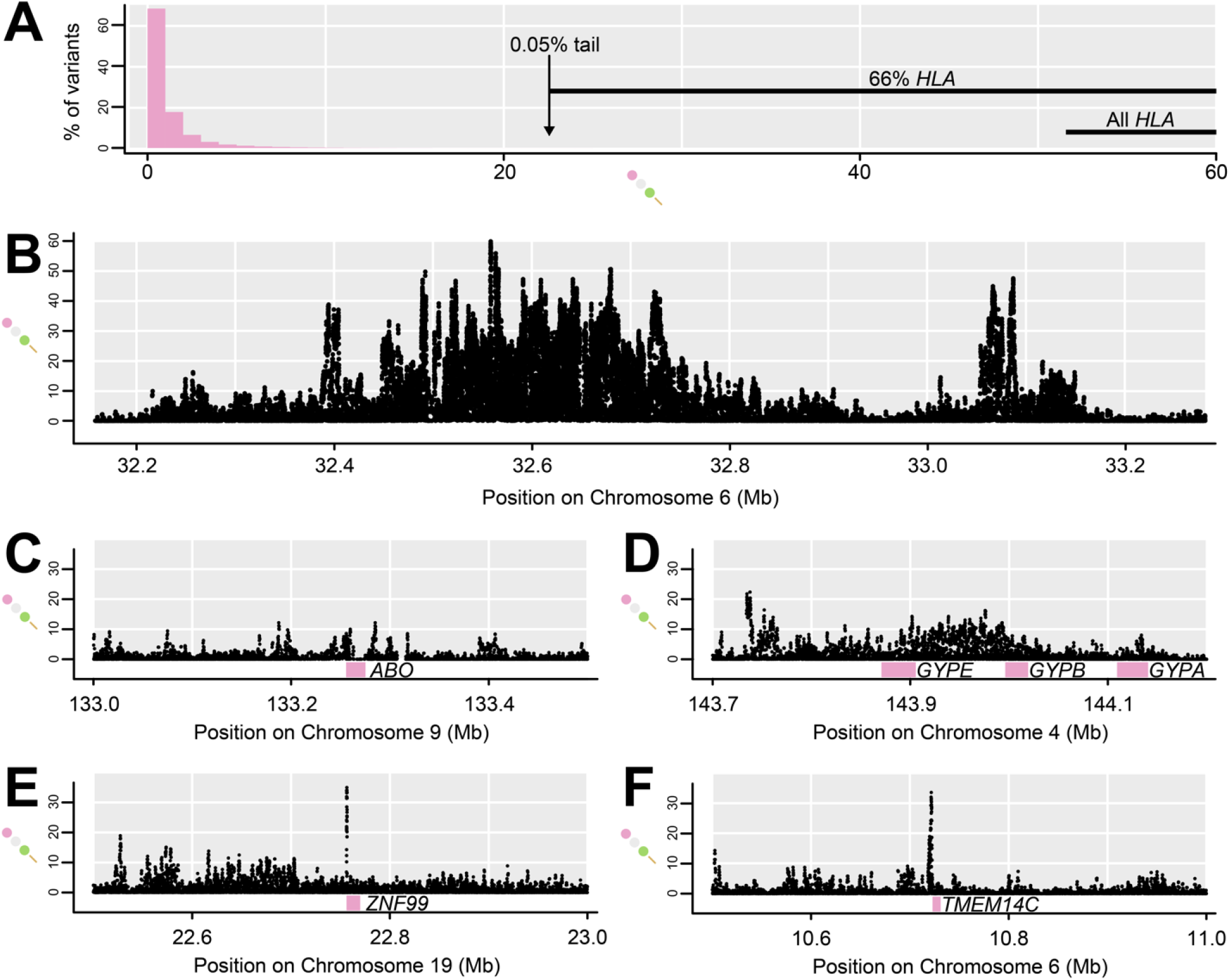
Distribution of 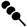, including notable genomic outliers. (A) Histogram of 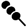 shows that the vast majority of variants have low values, and the 0.05% which exceed 22.6 are enriched for variants in or near the HLA. (B) This section of the HLA region shows the highest 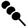 values in the genome. (C) *ABO* slightly surpasses 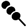 of 10 within the coding region. (D) Noncoding regions near glycophorin genes show high 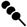. (E) The highest 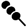 in or near a gene occurs just upstream of *ZNF99* (Table 2). (F) One of the highest 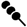 sites in or near a gene occurs just upstream of *TMEM14C* (Table 2).

### 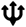 in simulations

Among 10,000 neutral simulated windows, the distribution of 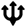 approximates a uniform distribution of expected p-values (Supp Figure 2). The fit is poor for higher values of 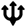, but since the practical question is whether low 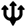 values are lower than would be expected by chance, this is unimportant. Upon adding a single window showing parallel adaptation to this set of 10,000, the adaptive window typically shows the lowest 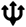. With a selection coefficient of 0.05, most (65%) adaptive windows have lower 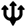 than all neutral windows, and 45% are individually significant 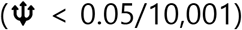, even though only 4% show F_ST_ higher than all neutral F_ST_ values for all three pairwise comparisons. Therefore, combining F_ST_ values into 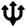 provides higher power to detect adaptation than individual pairwise F_ST_. With a stronger selection coefficient of 0.5, either 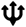 or F_ST_ alone perfectly distinguishes selection from neutrality.

### 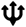 in human population data

In populations from Africa, Europe, and Asia, 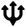 with *g* of 5 kb could be estimated for over 520,000 windows, representing over 2.6 Gb. The distribution of 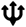 closely approximated a uniform neutral distribution of p-values, with 7% of windows showing 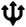 less than 0.05. The median number of common variants (MAF ≥5%) in these low-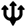 windows was 16, similar to the genome-wide median of 14. Only a single window, overlapping the majority of the coding sequence of gene *PTK6*, had an individually significant 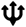 (less than corrected α of 1e-07; Figure 3A). However, windows with low 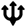 are good candidates for repeated shifts, even if not individually significant. Windows overlapping *ACKR1* occurred in the 0.03% most extreme windows 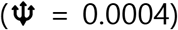, while windows overlapping *CR1* occurred in the 0.5% most extreme windows 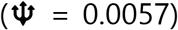. Low-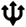 outliers are enriched for genic and exonic windows (Figure 3B), consistent with 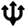 capturing adaptive variation. After staggering window starting positions to capture all outliers, the top 50 genes have an exon overlapping at least one window with 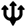 under 0.0002 (Supp Table 2). Results were largely similar with *g* of 50 kb (Figure 3C,D), indicating that 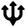 is robust to the choice of window size. Because these wider windows often overlap more than one gene, complicating interpretation, we focus on results with *g* of 5 kb (Table 2).

**Figure 3.**
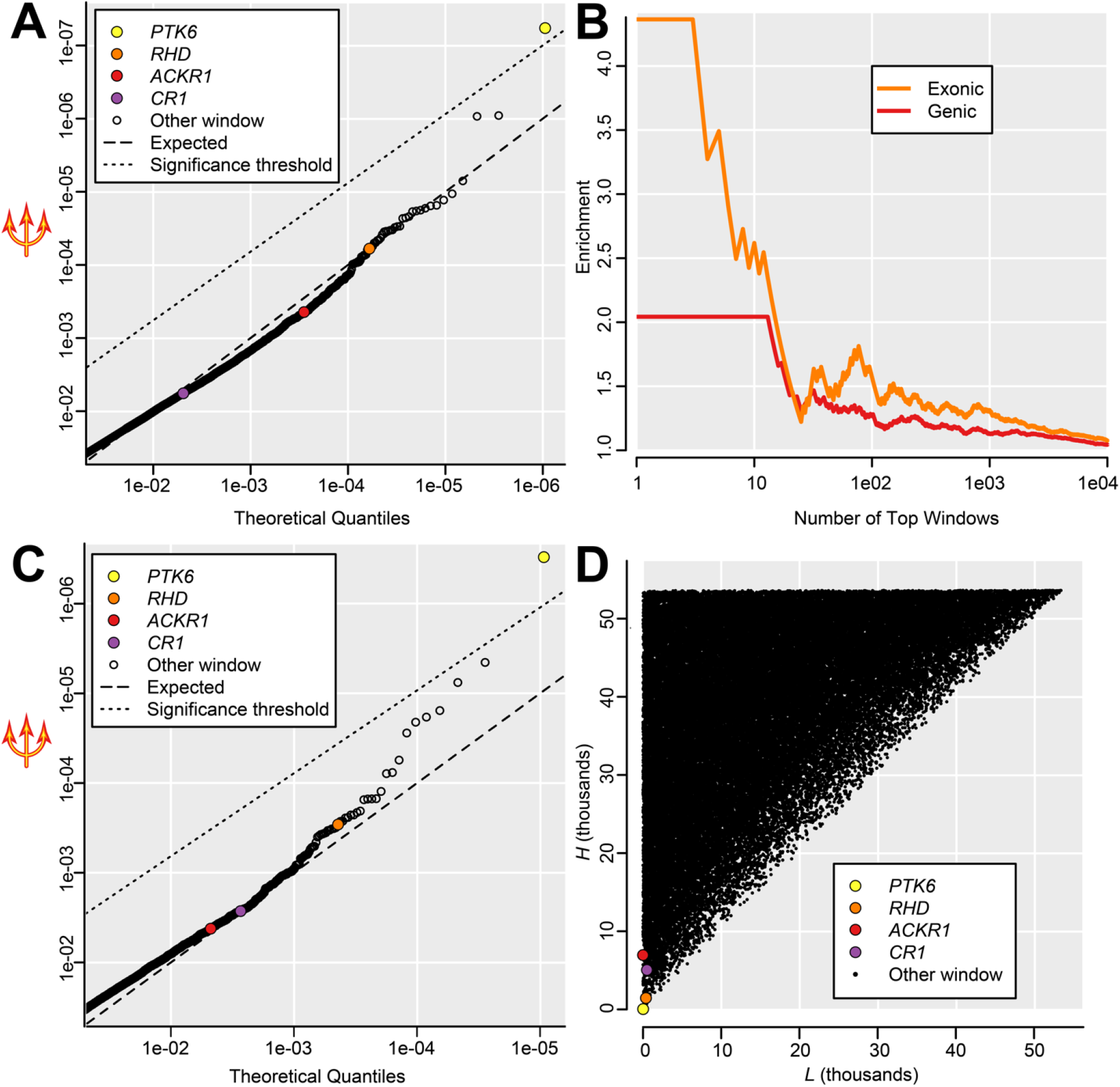
Genomic outliers for 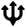. (A) Partial QQ plot showing 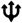 less than 0.05, representing 7% of genomic windows, for *g* = 5 kb. A single window overlapping *PTK6* surpasses the significance threshold, while most other windows are close to the expected neutral distribution. Although windows overlapping *ACKR1, CR1,* and *RHD* are not individually significant, they occur among the 0.5% most extreme windows. (B) 5 kb windows with low 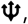 shown for continuously increasing thresholds, are enriched for windows that overlap genes, and even more so for windows that overlap exons, consistent with 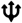 capturing phenotypically relevant polymorphism. (C) Partial QQ plot showing 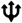 less than 0.05, representing 8% of genomic windows, for *g* = 50 kb. As in (A), a window overlapping *PTK6* is individually significant while *ACKR1, CR1,* and *RHD* are among the top outliers. (D) Lowest rank *L* (most extreme F_ST_) and highest rank *H* (least extreme F_ST_) for *g* = 50 kb, highlighting genes in outlier windows.

### 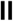 in simulations

In simulations, 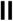 is substantially higher under an Arrested Sweep than under neutrality (Supp Figure 3). Under strong selection (heterozygote fitness = 1.99), almost all simulations produced higher 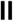 (median = 150; 95% interval = 64 to 237) than neutral simulations (median = 1; 95% interval = 0.01 to 32). Under weak selection (heterozygote fitness = 1.09), 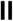 still typically exceeded the neutral distribution (median = 43; 95% interval = 17 to 78). 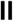 is calculated from several unrelated metrics, and each component has a distribution under selection that differs from the neutral distribution, resulting in a statistic that is very sensitive to Arrested Sweeps. In practice, these results suggest that under similar parameters, variants with 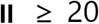 should be enriched for true balanced polymorphisms, while variants with 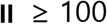 are very unlikely to be neutral.

### 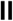 in human population data

Within our focal population YRI, using all five sub-Saharan African populations to calculate F_ST_ and *p*, and Europe as the outgroup, 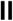 ranged from 0 to 511.2 (median = 1.0, 95% interval = 0.0 to 23.8; Figure 4A). The HLA region accounted for 4% of autosomal variants in the top 0.05%, with additional outliers closely linked to it including the top autosomal variant in an intron of *BAK1* (Figure 4B), but this HLA enrichment was much less pronounced for 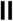 than it was for 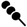. Our target genes *HBB* and *G6PD*, and specifically their phenotype-associated nonsynonymous polymorphisms, showed very high 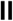 and are among the most extreme outliers. In *HBB*, the Glu-Val missense variant rs334 that causes sickle-cell anemia shows 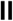 of 194.8, placing it in the top 0.005% of all variants (Figure 4C). Only 144 variants in the entire genome have a higher 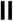 than rs334. If variants on chromosomes 6 (HLA) and X (see below) are ignored, rs334 remains among the top 14 variants, and the only one within a protein-coding gene. In *G6PD*, the Asn-Asp missense variant rs1050829 associated with G6PD deficiency shows 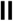 of 68.8, placing it in the top 0.5% of all variants, both on the X chromosome and genome-wide (Figure 4D). The top 50 genes outside of the HLA region (Supp Table 3; Figure 4C,E,F), based on 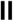 in exons or within 1 kb upstream, all exceed 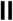 of 99.5 (top 0.025% of variants). This list includes several genes on chromosome 6 that could reflect the effect of HLA selection, as its signal appears to extend for several megabases surrounding the HLA (Figure 4B). While *HBB* is the highest autosomal gene, it is exceeded more than 2.5-fold by two adjacent X-linked genes, *ABCB7* and *RLIM* (Figure 4E). The *ABCB7* signal includes Ala-Val missense variant rs1340989 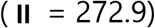 and intron variant rs372972791 with the highest observed 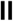 of 511.2. *ABCB7* is the peak of a 3 Mb region from X positions 74.5 to 77.5 Mb, with more than 100 variants showing 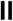 over 200, a threshold that excludes all other X-linked variants and all but 17 autosomal variants.

**Figure 4.**
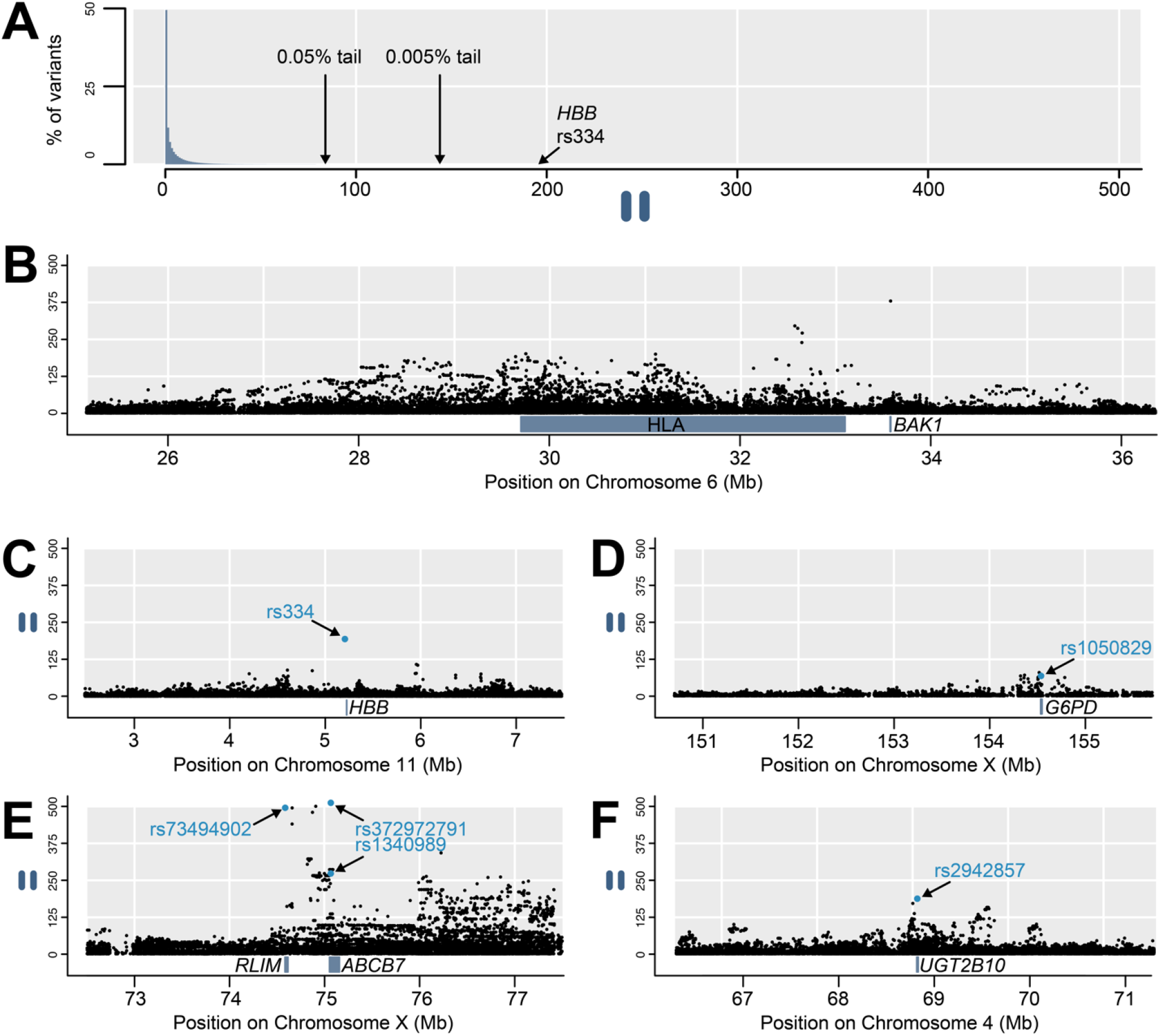
Distribution of 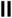, including notable genomic outliers. (A) Histogram of 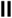 shows that the vast majority of variants have low values, though the sickle-cell polymorphism rs334 occurs in the 0.005% tail. High 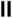 occurs throughout the HLA region and this signal extends beyond its borders, including in *BAK1*. Sickle-cell polymorphism rs334 in *HBB* shows exceptionally high 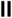. (D) Several variants in or near *G6PD* show high 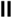, including disease-linked missense variant rs1050829. (E) The highest 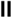 values in the genome by far occur in this section of the X chromosome, with the highest variants in *RLIM* and *ABCB7*. (F) Splice-acceptor variant rs2942857 in *UGT2B10* is the highest exonic autosomal variant after rs334.

### Synthesis

Among the top 50 outliers for each test, there is significant enrichment for the “antigen binding” molecular function GO term (9 genes, of which 8 are 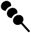 outliers, FDR = 0.049), and the “endocytosis” biological process GO term (16 genes, FDR = 0.038). We specifically tested for genes important to red blood cells, given their centrality to our exemplar genes and to *Plasmodium* invasion, and we observe substantial enrichment. Genes with a proteomic presence in erythrocytes are abundant, but not significantly so (1.3-fold enrichment; 23 genes; p> 0.05; Bryk and Wiśniewski 2017). There is significant enrichment for erythroid and erythrocyte genes as detected by text mining (Santos et al. 2015; Rouillard et al. 2016), both among the top 50 outliers for each test (1.8-fold enrichment; 17 of 150 genes; p < 0.05; Supp Tables 1, 2, and 3; Figure 5A), and among the top five outliers for each test (6.4-fold enrichment; 6 of 15 genes; p < 0.001; Table 2; Figure 5B). Enrichment is even greater among genes with higher text mining scores, for which there is stronger evidence for importance in red blood cells (Figure 5). Independent of this analysis, our top hits are significantly enriched for correlations with malaria susceptibility in genome-wide association studies (“GWAS”; 2.9-fold enrichment; 9 of 150 genes; p < 0.01; Supp Tables 1, 2, and 3). These nine GWAS hits include known exemplar gene *HBB* and adjacent gene pair *METTL7B* and *ITGA7* that share a signal with each other, but even if *HBB* is discarded and the adjacent pair is merged, the enrichment is still significant for seven matches (p < 0.05). We do not see an enrichment for proteins previously shown to interact with *Plasmodium* or piroplasmid parasites (p> 0.05). Many genes are observed both among our outliers and among those of nine previous genome-wide scans for selection using various tests (Supp Table 4). There is a trend toward enrichment with all nine previous scans, though it’s not always significant. Notable comparisons include five genes identified by both 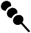 and a composite likelihood scan for balancing selection (DeGiorgio et al. 2014; 13.4-fold enrichment, p < 1e-04), thirteen genes identified by both 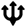 and the parallel adaptive divergence scan at the level of individual variants (Tennessen and Akey 2011; 4.1-fold enrichment; p < 1e-05), and two genes identified by both 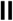 and iHS (Voight et al. 2006; 2.9-fold enrichment; p> 0.05). Finally, across all tests we see an enrichment for low-recombination regions of the genome (2.5-fold enrichment; 20 of 150 genes; p = 0.0001), especially for 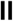 which encompasses long-range LD (Table 2; Supp Tables 1, 2, and 3). We did not exclude these low-recombination genes, because we suspect they are not false positives but rather reflect enhanced power to detect selection in these regions, and because our 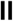 exemplar gene *G6PD* occurs in a low-recombination region.

**Figure 5.**
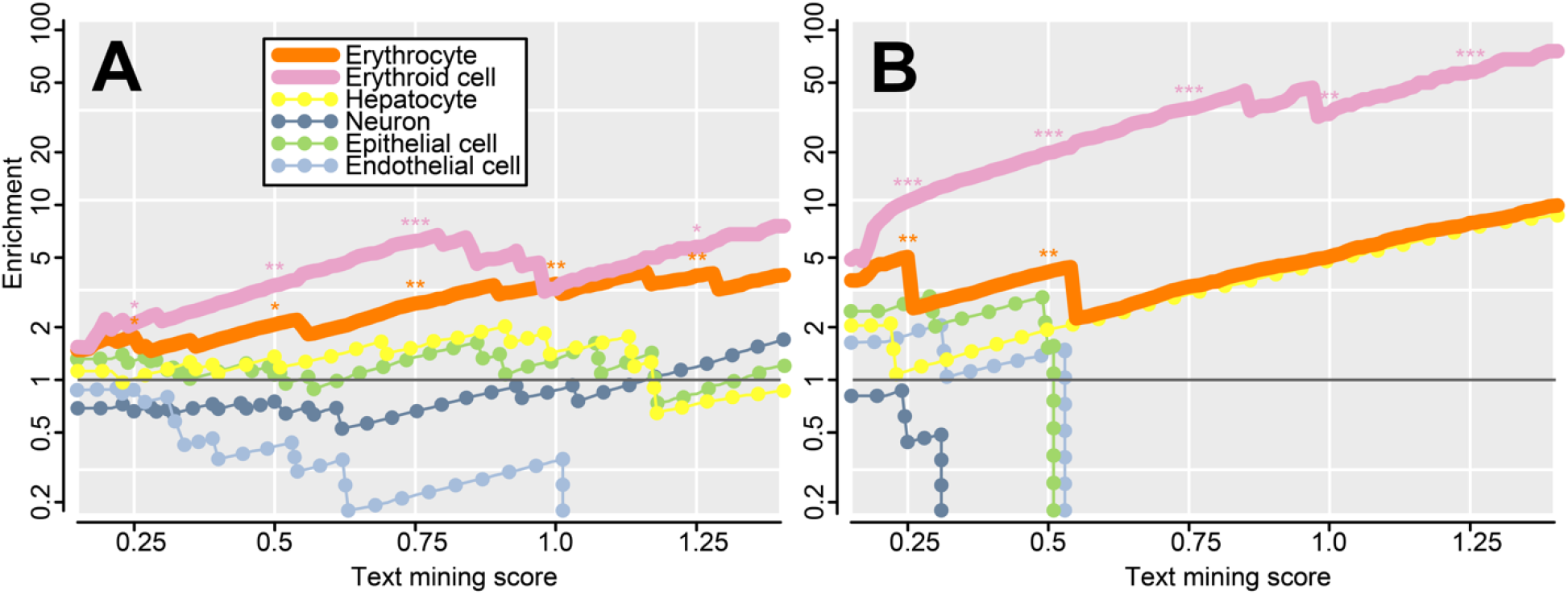
Enrichment for genes associated with various cell types, with higher scores indicating a stronger association via text mining (Santos et al. 2015; Rouillard et al. 2016). At intervals of 0.25, significant enrichments are indicated (*p < 0.05; **p < 0.01; ***p<0.001). Genes associated with erythrocytes and erythroid cells are significantly enriched (solid lines), and enrichment increases with score, suggesting a prevalence of genes with particularly high specificity to red blood cells. Enrichment is higher among erythroid cell genes than among erythrocyte genes, suggesting that many outliers are more important in erythroid progenitors than in mature cells. No significant enrichment is observed in control tissues (dotted lines). (A) The top 150 genes for the three tests (Supp Tables 1, 2, and 3). (B) The top fifteen genes for the three tests (Table 2).

## Discussion

### Three new statistical assays for non-neutral polymorphism

We present three novel summary statistics that reflect evidence for selective neutrality in population genetic data. There are now many such statistical tests (Vitti et al. 2013), which are frequently employed to find putative targets of selection across the human genome (Sabeti et al. 2007; Akey 2009; Fan et al. 2016). However, despite this plethora of statistical tools, it remains challenging to conclusively identify instances of positive or balancing selection in humans. Genes which are known to behave non-neutrally, either because fitness or phenotypic impact of genotypes have been measured directly (e.g. overdominance at *HBB*; Aidoo et al. 2002), or because of evidence from other species (e.g. trans-species polymorphisms at *ABO* and *GYPA/B/E;* Ségurel et al. 2012; Leffler et al. 2013), are not necessarily outliers in scans for selection across human populations. In the approach presented here, we focused on genes with polymorphisms already known to be adaptive and shaped by infectious disease, and we attempted to capture unusual empirical patterns at these genes that are plausibly driven by selection in accordance with population genetic theory. Our statistics are inspired by patterns at malaria-relevant genes in humans and they reveal numerous outliers with potential relevance to malaria. This is because malaria has been such a strong selective pressure on our species, and because our adaptive response to *Plasmodium* frequently leads to these population genetic patterns, making them likely indicators of malaria resistance loci. However, malaria is not the only selective pressure capable of producing these patterns, and these tests are potentially applicable to detecting selection due to other causes and/or in other species.

Both simulated and empirical results suggest that these tests are robust. However, they carry several caveats and limitations. Like other outlier-based tests (Akey 2009), these tests are intended to identify genomic regions that are most suggestive of certain hypothesized modes of selection, under the *a priori* assumption that a small proportion of the genome did evolve according these modes. They are not intended to test a null hypothesis of complete neutrality, and there are no clearly defined thresholds beyond which neutrality can be rejected. None of the tests employ outgroup species and so are naïve with respect to derived or ancestral status of alleles, but if selective pressures fluctuate and act on standing variation then even an ancestral allele could be adaptive. As with most selection scans, these tests assume that mutation and recombination rates across the genome are similar; as this assumption is typically violated, their power may vary across genomic regions. Each test also carries its own specific caveats. For Divergent Haplotypes, selectively neutral processes like gene conversion or introgression (e.g. from archaic hominids; Ragsdale and Gravel 2019) could lead to high sequence divergence between haplotypes and thus elevated 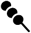, as could mis-aligned reads from paralogs. Furthermore, the most polymorphic genomic regions can be poorly represented in population genomic datasets if high sequence divergence and large indels impede genotyping, so some of the strongest Divergent Haplotypes signatures could be missed by 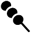 in practice. For Repeated Shifts, 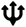 approximates a p-value but is not formally a p-value for a defined null hypothesis. F_ST_ depends on the MAF, and single-variant-based tests for parallel F_ST_ require considerable filtering based on allele frequency (Tennessen and Akey 2011). However, since 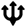 considers F_ST_ across numerous variants within a window, typically showing a wide range of MAFs, it should be largely robust to this effect. For Arrested Sweeps, 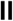 carries the assumption that differences in heterozygosity are caused by a shared physical and evolutionary association with the target locus. This assumption is violated if there is LD not caused by physical linkage (i.e. due to family or population structure), or if LD due to sampling error is substantial enough to affect the true LD signal. The latter can occur when regions of extraordinarily high polymorphism or haplotype structure such as a centromere or the HLA (Figure 4B) are included in the window used to estimate heterozygosity. 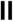 depends on selection acting consistently one way in the ingroup and another way in the outgroup, which may not always be the case. All of these statistics can best be thought of as tools for identifying candidate genes, but follow-up study is required before drawing firm conclusions about evolutionary history or functional impact.

The tests detect the six exemplar loci which motivated this study, but with varying degrees of success. Most strikingly, 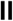 is a near-perfect test for an *HBB*-like signal, as *HBB* is the third most extreme gene and the top autosomal gene. *ACKR1* and *G6PD* are also notable outliers, falling in the top 1% of genes for their respective statistics. The remaining three exemplar loci are less extreme outliers, but all fall within the top 5% of genes for their respective statistics. As noted above, 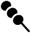 may be underpowered if some polymorphisms are absent from the dataset, as is the case for *ABO and GYPA/B/E*, which could partially explain why this statistic was the least powerful at detecting its exemplar loci. Other loci are also strongly associated with malaria but would have made poor exemplar genes in this analysis and were therefore ignored. For example, *Plasmodium* has driven an Arrested Sweep on *SLC4A1* in Southeast Asia (Paquette et al. 2015), but these populations are poorly represented in the 1000 Genomes. Also, though *ATP2B4* is globally associated with malaria (Malaria Genomic Epidemiology Network 2019), evidence for non-neutral evolution is mild and/or very geographically localized (Gelabert et al. 2017; Gouveia et al. 2019). Indeed, *ATP2B4* was not a notable outlier in any test (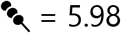, 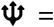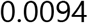, 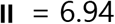).

### Hallmarks of adaptation to malaria

Malaria is caused by *Plasmodium*, which for millions of years has been a parasite of hominid primates and other vertebrates, yielding the three hallmarks of selection presented here (Table 1). Each hallmark has occurred on a different timescale and is associated with distinct parasite pressures. First, genetic variants which conveyed resistance to ancient *Plasmodium* parasites in our pre-human ancestors, if not fixed long ago, persist as balanced polymorphisms with a Divergent Haplotypes signature. These variants, at loci like *ABO, GYPA/B/E*, and the HLA, have also been shaped by other pathogens over millions of years. The importance of *Plasmodium* itself as a driver of diversity is unknown but highly plausible given the connection between these loci and malaria resistance (Ségurel et al. 2013). Second, the Repeated Shifts pattern is tied to the emergence of modern *Plasmodium* species from parasites of non-human apes, exerting novel selection pressures on early modern humans. Evidence suggests that *P. vivax* originated first, over 20,000 years ago, while *P. falciparum* only arose within the past 10,000 years (Loy et al. 2017; Daron et al. 2020). The dramatic selective sweep of the Duffy-null allele in Africa conveyed resistance to *P. vivax* and could have eliminated it as a human parasite, but a lineage of *P. vivax* survived in Asia where it persists widely to this day (Hamblin and Di Rienzo 2000; Loy et al. 2017). It is unclear why Duffy-null did not also spread widely beyond Africa, but instead *P. vivax* in Asia selected for a different *ACKR1* allele, Fy^a^ (King et al. 2011; Chittoria et al. 2012). *CR1* shows a similar signature of parallel evolution and impacts resistance to both *P. vivax* (Prajapati et al. 2019) and *P. falciparum* (Tham et al. 2010), suggesting that either or both of these species may have driven its evolution. Third, Arrested Sweeps are the most recent evolutionary scenario and are associated with the expansion of *P. falciparum,* which likely dispersed across sub-Saharan Africa along with agriculture, which facilitates optimal mosquito habitat, during the past few thousand years. Resistance alleles at *HBB* and *G6PD* spread among the same locations on a similar timescale (Tishkoff et al. 2001; Shriner and Rotimi 2018; Laval et al. 2019). Each hallmark of selection might only be detectable within its particular timescale. For example, an Arrested Sweep polymorphism maintained by a deleterious homozygote genotype is inherently unstable and poised to be replaced by an allele that conveys the heterozygote advantage without the costs, as has already begun for *HBB* (Modiano et al. 2001). Thus, any hypothetical Arrested Sweep caused by *P. vivax* prior to the Duffy-null sweep could have been subsequently lost.

We observed an enrichment for malaria-associated chromosomal regions from three large GWAS (Timmann et al. 2012; Malaria Genomic Epidemiology Network 2019; Milet et al. 2019), especially among 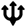 outliers (Supp Table 2). Two particularly promising candidates are *PTPRM* and *MYLK4*, which are among the very strongest candidates for recurrence of mild malaria attacks in infants (Milet et al. 2019). In particular, *PTPRM* was the top prioritized gene in the GWAS based on functional consequences (p = 3.8e-08; Milet et al. 2019), and the top disease-associated variant in the gene occurs in between the two variants that define the Repeated Shifts signal 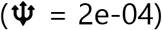, just upstream of an alternate transcript start. Furthermore, less than 2 kb upstream of malaria-associated *SPNS2,* the fourth-highest 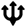 outlier (Table 2), there are three variants within 36 bp of each other that have undergone three distinct allele frequency shifts in Africa, Asia, and Europe. Finally, adjacent genes *METTL7B* and *ITGA7* share a Repeated Shifts signal and a GWAS signal and are both overexpressed during severe malaria (Lee et al. 2018). Any gene at the intersect of selection signal and phenotype association is worthy of further consideration, as such combined evidence has been instrumental in implicating known loci like *GYPA/B/E* (Malaria Genomic Epidemiology Network 2015).

### Erythrocytes and erythropoiesis

Of particular interest with respect to malaria is the pronounced enrichment for genes with a role in red blood cells (Figure 5). The most severe pathology impacting human fitness occurs during the blood stage of the *Plasmodium* life cycle, and our exemplar genes on which we based our tests all act during this stage. There are two principal mechanisms by which such genes may impact blood-borne parasites. In the first mechanism, erythrocyte surface proteins encoded by transmembrane genes like *ACKR1*, *CR1*, *ABO*, *GYPA*, and *GYPB* act as receptors for *Plasmodium* ligands to facilitate invasion (Cowman et al. 2017) or mediate cytoadherence (Cserti and Dzik 2007). Several such receptors remain to be discovered (Cowman et al. 2017), but there are few compelling candidates among our novel outliers. The strongest contender is *RHD* (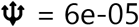; Figure 3), which encodes the transmembrane D antigen for the Rh blood group, a component of the erythrocyte cell surface connected to *Plasmodium* invasion (Chung et al 2008). Remarkably, *RHD* also shows a moderate Divergent Haplotypes signature within its coding region (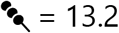, higher than either of the Divergent Haplotypes exemplar genes), suggesting it could have been subject to both adaptive processes. While the signal of selection on *RHD* is convincing, unlike *ACKR1* and *CR1* this signal involves Europe-specific divergence and thus shows a slightly different evolutionary history, perhaps driven by other parasites (Novotná et al. 2008). Many other outliers also encode transmembrane proteins, but there is little evidence that they are expressed on the surface of mature erythrocytes.

The second principal mechanism of resistance is to alter erythrocyte development and cellular integrity. Such changes can affect intracellular parasite growth and survival, though perhaps with reduced function or a similar cost to the host. Hemoglobinopathies and enzymopathies that protect against malaria are conveyed by *HBB*, *G6PD*, and other loci like *FECH* (Taylor et al. 2013; Smith et al. 2015). Across all three tests, several of our top outlier loci play keys roles in erythropoiesis (Table 2; Supp Tables 1, 2, and 3), and thus we observe more evidence for adaptation via this mechanism than via the first mechanism of invasion receptors. Loci involved in erythroid development are not necessarily expressed in the mature proteome (Bryk and Wiśniewski 2017), and thus we see greater enrichment for (precursor) erythroid cell genes than for (mature) erythrocyte genes (Figure 5). In addition to *HBB*, our most extreme outliers include *TMEM14C* and *ABCB7,* both implicated in erythroid maturation (Conte et al. 2015; Table 2). *TMEM14C* encodes a transmembrane protein essential for erythroid synthesis of heme (Yien et al. 2014), and many of the highest 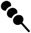 variants occur in a 4.5 kb region in its upstream cis-regulatory region (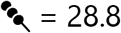, Figure 2F). As further evidence for malaria-relevant selection independent of the 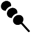 signal, variants near *TMEM14C* are also among the most differentiated between Europeans and Africans genome-wide (F_ST_> 0.8) and may underlie the pronounced differences in *TMEM14C* expression between these continents (Quach et al. 2016). *ABCB7*, a transmembrane iron transporter in the heme pathway, is essential for erythropoiesis and causes anemia when deficient (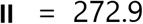; Pondarre et al. 2007; Severance and Hamza 2009; Figure 4E). The outlier region centered on *ABCB7/RLIM* far exceeds 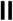 in the rest of the genome. Recombination is unusually low in this region, which may partially explain the magnitude of the signal, but low recombination alone should not produce a 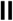 outlier in the absence of selection. Like *TMEM14C* and *ABCB7*, several outliers are directly involved in heme binding and/or biosynthesis (*PRDX1, UROD, CYB5R3*; Fermo et al. 2008; Severance and Hamza 2009), while liver-expressed *UGT2B10* is part of a glycosyltransferase family that catalyzes heme breakdown (Sticova and Jirsa 2013; Figure 4F). Other outliers have roles in hematopoiesis (*RABGAP1L*, Roberti et al. 2009; *MAP1LC3B*, Kang et al. 2012) or erythrocyte morphology (*MYH9*, Smith et al. 2019).

### Adaptation to infectious disease beyond malaria and the red cell

Malaria is only one important selective pressure in humans, along with climate, diet, environmental toxins, and other infectious diseases (Fan et al. 2016). Consistent with these expectations, many of the outliers in our tests have no obvious connection to *Plasmodium*. Our lists of top outliers (Supp Tables 1, 2, and 3) do not closely match any particular previous scan for selection, though there is enrichment for repeat outliers (Supp Table 4), including *LGALS8* (Andrés et al. 2009; DeGiorgio et al. 2014; Bitarello et al. 2018), *FBXO31* (Leffler et al. 2013), *SORD* (Tennessen and Akey 2011; DeGiorgio et al. 2014), and *DMBT1* (Leffler et al. 2013; DeGiorgio et al. 2014; Siewert and Voight 2017). One of the clearest signals of selection is on *PTK6* (Figure 3), a tyrosine-protein kinase involved in several cancer pathways. *PTK6* also shows parallel evolution at the variant level (Tennessen and Akey 2011) and is speculated to harbor adaptive polymorphisms impacting gastric bacterial infection (Jha et al. 2015). However, the specific selective pressure on *PTK6*, and most other outliers, is unknown.

Our outliers include numerous immune-related genes, which are especially prone to positive and balancing selection (Barreiro and Quintana-Murci 2010; Spurgin and Richardson 2010). This trend does not preclude a role for *Plasmodium*, but it is also consistent with selection via other infectious agents from viruses to macroparasites. Many HLA-linked variants are outliers for 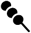 and 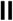, but we have largely ignored these, as balancing selection on the HLA is already well documented (Spurgin and Richardson 2010). Excluding them, the 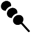 outliers are enriched for “positive regulation of immune response” and “antigen-binding”, a trend largely driven by immunoglobulin genes (Supp Table 1). Cumulatively across all three tests there is enrichment for “antigen-binding” and “endocytosis” which also encompasses the immunoglobulin genes as well as other immune-relevant genes like dendritic cell receptor *CD209*, T-cell surface glycoprotein *CD5*, and macrophage-expressed *CLEC4F*. Many of the immunity-related outliers encode transmembrane proteins, especially outliers for 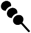 and by a lesser extent 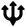. This pattern is consistent with selection for novelty in parasite-recognition proteins, leading to stable negative frequency-dependent selection (Divergent Haplotypes) or regular positive selection for new variants (Repeated Shifts). In contrast, for 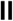 neither the exemplar loci (*HBB* and *G6PD*) nor most of the empirical outliers (Supp Table 3) encode transmembrane proteins, and they show fewer direct ties to the immune system. Therefore, to the extent that Arrested Sweeps reflect selection by infectious agents, the adaptive response appears to compromise basic metabolic cytoplasmic proteins and thus prevent pathogens from rising to overwhelming levels, though perhaps at a cost to the host.

### Future directions

The main goal of this study was to develop and evaluate statistical metrics for detecting malaria-associated signatures of selection. We used the 1000 Genomes as a reliable and comprehensive dataset for this purpose, but future work on additional emerging datasets could further clarify patterns of selection (e.g. GenomeAsia100K Consortium 2019). These tests may be underpowered here due to variants being absent from the dataset, including large indels and copy-number variants, but this issue can be addressed with more complete, high-coverage sets of genotypes. Furthermore, these methods are valid to apply to other species to detect signals of selection driven either by infection or by other factors. Scripts which calculate the statistics presented here are available at https://github.com/jacobtennessen/MalariaHallmarks. The identification and validation of additional examples of functional adaptive polymorphism will allow further refinement of tests for selection, leading to even more discoveries in a fortuitous feedback loop. In this way, the fields of evolutionary genetics and malaria pathology will continue to bolster each other as they have done for decades.

## Materials and Methods

### Divergent Haplotypes and 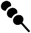

For any pair of loci *i* and *j* with MAFs *p_i_* and *p_j_* and joint minor frequency *p_ij_*, one measure of LD between them (Kelly 1997) is defined as:

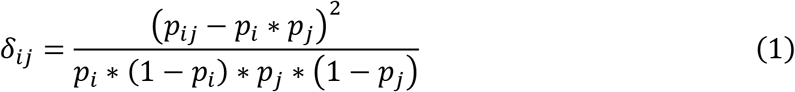

The mean LD for a set of *S* adjacent variants in *n* sequences (Kelly 1997) is thus:

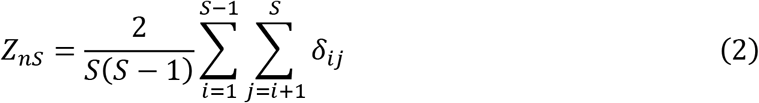

For a target variant *j* and the *S* - 1 other variants *i* within distance *g* bp of *j*, in *n* sequences, the statistic D_ng_ or 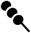 (“Dango”) is defined here as:

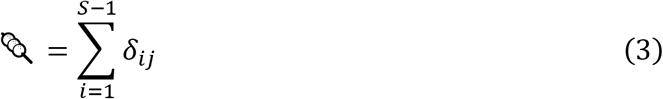

Because LD can be affected by population structure, 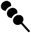 should be calculated for individual populations consistent with panmixia, and not across populations that differ in allele frequencies.

### Simulations with 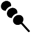

In order to evaluate 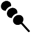, we used the forward-time evolution simulation package SLiM (Messer 2013). We simulated genomic windows of 10,001 bp, with an overdominant balanced polymorphism in the center. 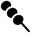 is not expected to only detect overdominance, which is only one type of balancing selection, but overdominance is logistically straightforward to simulate. By default, the dominance coefficient was 1e+06 and the selection coefficient was 1e-08, yielding nearly identical homozygote finesses of effectively 1, a heterozygote fitness of 1.01 (= 1+ 1e+06*1e-08), and an expected MAF of 0.5. We also considered a “skew” scenario with uneven fitnesses: a dominance coefficient of 1.1 and a selection coefficient of 0.1, yielding homozygote fitnesses of 1 and 1.1, a heterozygote fitness of 1.11, and an expected MAF of 0.08. All other polymorphisms were selectively neutral and generated with mutation rate (μ) of 1e-7. We simulated either 50,000, 100,000, or 200,000 generation of evolution in a population of 10,000 individuals, with population-scaled recombination rate (ρ) set to either 0.01 or 0.001. We then calculated 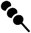 for the balanced polymorphism, using *g* ranging from 100 to 5000 bp. As a control, we simulated windows in which all polymorphisms were selectively neutral. These neutral control windows were 15,000 bp, and from each we randomly chose a single variant for which to calculate 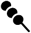, with MAF ≥ 0.4 and at least 5,000 bp of sequence on either side. Other parameters matched the selection simulations, with one addition: we also considered a scenario with neutral evolution but a doubled μ of 2e-7, to test if elevated mutation rate alone can be distinguished from a signal of selection. For each distinct set of parameters, we ran 1000 replicate simulations. We quantified overlap between the distributions of simulated windows by finding the lower 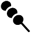 quantile in selection simulations that matched the equivalent upper quantile of neutral simulations.

### Application of 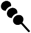

We scanned for 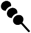 in YRI in the 1000 Genomes dataset (1000 Genomes Project Consortium 2015), using a distance of *g* = 500 bp. We defined the top 50 candidate genes by ranking all protein-coding genes occurring outside of the HLA region (chromosome 6 between 29.7 and 33.1 Mb) based on 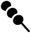 within exons or within 1 kb upstream of the gene, under the assumption that these sections are the mostly likely to harbor functional polymorphisms.

### Repeated Shifts and 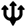

Consider a set of three populations. For *R* nonoverlapping genomic windows of size *g* bp, excluding any windows with fewer than two variants, one calculates the highest F_ST_ among all variants, for each of the three pairwise comparisons. For each pairwise comparison, one then ranks all windows by F_ST_, using integers from 1 (highest F_ST_) to *R* (lowest F_ST_), such that higher F_ST_ values are ranked lower. Ties are rounded up; e.g. if the highest F_ST_ value for a given population pair is observed in two different windows, both windows are assigned a rank of 2 and no window is assigned a rank of 1. Each window thus has three ranks, one for each population pair. For each window, the lowest rank *L* (most extreme F_ST_) and the highest rank *H* (least extreme F_ST_) are then identified. The statistic T_R_ or 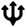 (“Trident”) is defined as:

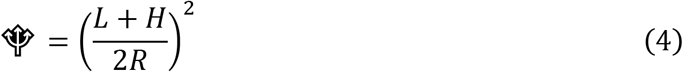

### Simulations with 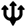

As with Divergent Haplotypes, we performed forward-time simulations using SLiM. We simulated 10,000 unlinked genomic windows of 5000 bp, with μ 1e-7, recombination rate 0 within windows, and all mutations selectively neutral. We allowed a population of 10,000 diploid individuals to evolve for 14,000 generations, at which point a second population of 5000 individuals is generated from the first one. At 14,500 generations a third population of 5000 individuals is generated from the second one, and all three populations continue to evolve for 500 more generations until the 15,000th generation. We then calculated F_ST_ values by randomly sampling 500 individuals per population. Furthermore, we simulated additional windows under the same parameters but with two adaptive mutations arising at generation 14,750: one in population 1 and one in population 3. For 1000 of these windows we used a selection coefficient of 0.05, and for 1000 of these windows we used a selection coefficient of 0.5. The adaptive mutations occur at different sites in the window and initially appear with 50 copies per population, to minimize the chance that they are lost to drift; this can be thought of as an existing rare neutral mutation suddenly becoming adaptive, or else a novel mutation occurring several generations earlier and reaching an abundance of 50 by generation 14,750. To calculate 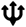, we combined each adaptive window with the 10,000 neutral windows one at a time, rather than including all adaptive windows together, to simulate a genome in which the vast majority of windows are neutral. Thus, we could simultaneously evaluate whether the neutral windows behaved neutrally and whether the single adaptive window appeared as an outlier.

### Application of 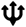

As with Divergent Haplotypes, we scanned the 1000 Genomes dataset for 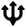 (1000 Genomes Project Consortium 2015). Our populations were Africa, Europe, and Asia. To maximize the signal of local adaptation and minimize admixture, we calculated African allele frequencies from all 504 individuals from the five sub-Saharan African populations (ESN, GWD, LWK, MSL, YRI) and ignored the two diaspora populations (ACB, ASW). For Europe we used all 503 individuals from the five populations (CEU, TSI, FIN, GBR, IBS), and for East Asia we used all 993 individuals from the ten South Asian and East Asian populations (CHB, JPT, CHS, CDX, KHV, GIH, PJL, BEB, STU, ITU). We used window sizes of *g* = 5000 bp and *g* = 50,000 bp. We only examined autosomes to avoid the confounding effects of the X chromosome’s unique evolutionary rate impacting F_ST_. By default, we aligned windows beginning at the start of each chromosome. However, because windows are nonoverlapping, 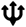 is sensitive to how windows are aligned; two closely adjacent variants could be assigned to different windows and thus their shared signal would be missed. Therefore, we also calculated 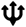 by starting windows at each 500 bp interval between 0 and 4500 bp from the start of each chromosome. We defined the top 50 candidate genes by ranking protein-coding genes according to top 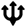 in windows overlapping exons.

### Arrested Sweeps and 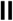

For a given variant with alleles A and a, one calculates the absolute difference in total heterozygous sites between individuals homozygous for allele A (*H_AA_*) and individuals homozygous for allele a (*H_aa_*). This difference can be quite large for rare variants, but these are uninteresting with respect to selection; instead, the signal of a sweep is a large difference for a variant that has risen beyond rarity (>1% frequency). Thus, the difference is multiplied by the MAF, which is calculated across the ingroup and designated *p_j_* as above. The product is the heterozygosity difference associated with allele A, *A_H_*, for which a high value indicates that one allele is associated with much more nucleotide diversity than the other, a signal of natural selection sweeping away variation in an otherwise polymorphic region:

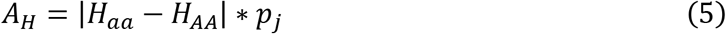

The distribution of F_ST_ in the ingroup will depend on *p_j_* and on the particular populations examined and can extend below zero under Weir and Cockerham’s (1984) formula, but all that matters it the relative, not absolute, value of F_ST_. Thus, one ranks all ingroup F_ST_ values for variants with MAF of *p_j_* (rounded to the nearest 1% in practice), with higher F_ST_ values getting lower ranks as with 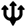. For each rounded *p_j_*:, ranks are divided by the total, yielding an F_ST_ rank proportion, *F_R_*, which ranges from 0 (high F_ST_) to 1 (low F_ST_, which is the relevant signal of selection in this scenario). The purifying selection metric is an adjusted reciprocal of MAF in the outgroup, *p_o_*, centered around a MAF of 1%. If *p_o_* is 0, the adjusted reciprocal is 1 and does not change the final product. If *p_o_* is 1%, the adjusted reciprocal is 0.5. As *p_o_* increases above 1%, the adjusted reciprocal rapidly declines, indicating low evidence for an Arrested Sweep. Thus, the measure of a variant showing similar frequencies in the ingroup while excluded from the outgroup, *z*, is:

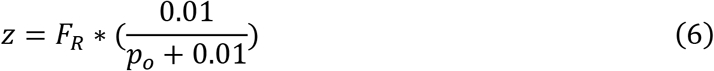

This product of these sweep and arrest metrics, Π_AHz_ or 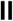 (“Pause”), is thus:

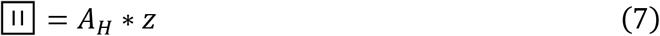

To calculate 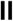 for a variant in a set of *n* phased diploid samples, one first calculates the total number of heterozygous sites for every possible diploid genome that could be formed from the 2*n* phased haploid genomes (2*n* choose 2 combinations). In practice, one can assume that the variant does not affect heterozygosity farther away than a given distance *g* (here set as 1 Mb) on the same chromosome, and thus heterozygosity can be calculated for a sufficiently large window on either side of the variant, rather than for the entire genome. This calculation can be performed once for a large genomic window (here set as 5 Mb) and then applied to all variants that are at least *g* from the edge of the window. For the target variant, one identifies all homozygotes for either allele among the 2*n*-choose-2 genomes, and calculates the mean number of heterozygous sites for each, yielding *H_aa_* and *H_AA_*. It is arbitrary which allele is designated as A versus a, and it does not depend on which is derived, dominant, etc. The absolute value of the difference between *H_aa_* and *H_AA_* is then multiplied by ingroup MAF *p_j_*, the F_ST_ rank *F_R_*, and the adjusted reciprocal of the outgroup MAF, (0.01/(*p_o_* + 0.01). As with 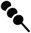, this test targets a single panmictic population in order to minimize the effect of population structure on LD.

### Simulations with 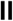

As with the other statistics, we performed forward-time simulations using SLiM. We simulated genomic windows of 2,000,001 bp, with an asymmetrical overdominant balanced polymorphism in the center. As with 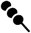, overdominance is not the only form of balancing selection that 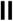 could detect, but it is the representative form used in simulations. We considered “strong” and “weak” selection scenarios. In the “strong” scenario, selection against derived homozygotes was −0.99 and the dominance coefficient was −1, yielding genotype finesses of 1 (ancestral homozygote), 1.99 (heterozygote), and 0.01 (derived homozygote), and an expected MAF of 0.33. In the “weak” scenario, selection against derived homozygotes was −0.9 and the dominance coefficient was −0.1, yielding genotype finesses of 1 (ancestral homozygote), 1.09 (heterozygote), and 0.1 (derived homozygote), and an expected MAF of 0.08. All other polymorphisms were selectively neutral and generated with μ of 1e-7. We set ρ to 0.001. We first simulated 10,000 generations of neutral evolution in a population of 10,000 individuals, then the outgroup population of 10,000 individuals was generated from the initial population. After another 2000 generations of neutral evolution, the adaptive mutation was generated in a single sample in the initial population. Unlike with 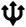, here it is important for the mutation to first appear in a single individual to generate the change in LD as the rare haplotype rapidly increases in frequency. After 100 additional generations, four new populations of 10,000 individuals were generated from the initial population to form the ingroup. The simulation then proceeded for 200 more generations to allow the ingroup populations to diverge. Thus, after a total of 12,300 generations we calculated 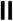 for the balanced polymorphism. As a control, we simulated windows in which all polymorphisms were selectively neutral. These neutral controls windows were 2,005,001 bp, and from each we randomly chose a single variant for which to calculate 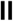, with MAF ≥ 0.05 and at least 1 Mb of sequence on either side. Other parameters matched the selection simulations. In some simulations the target polymorphism was lost to drift while rare, but these were subsequently ignored. For each of the three scenarios (strong selection, weak selection, and neutrality) we examined 1000 replicate simulations in which the target polymorphism was retained.

### Application of 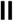

As with the other statistics, we scanned the 1000 Genomes data for 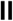 (1000 Genomes Project Consortium 2015). We again used YRI as our target population. We used the five sub-Saharan African populations (ESN, GWD, LWK, MSL, YRI) as the ingroup under the assumption that all have experienced similar disease-induced selection. We used Europe (CEU, TSI, FIN, GBR, IBS) as the outgroup. We assumed that only variation within *g* = 1 Mb was relevant to a given variant. We avoided all sequence within 5 Mb of the centromere on all chromosomes, because unusual levels of polymorphism and LD in these regions could swamp the signal. We only considered variants with *p_j_* of at least 0.01. We defined the top 50 candidate genes using the same criteria as for 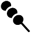.

### Synthesis

We identified significant gene ontology (GO) terms using false discovery rate (FDR) values from http://geneontology.org. For all other tests for enrichment, we used Fisher’s exact tests. We tested for enrichment in the erythrocyte proteome using the genes detected by Bryk and Wiśniewski (2017). We also compared our top outliers against “erythroid cell” and “erythrocyte” genes on Harmonizome (http://amp.pharm.mssm.edu/Harmonizome; Rouillard et al. 2016), a compilation of text-mining databases of genes and tissues (Santos et al. 2015). To formally test for enrichment, we included the 507 erythroid genes and 1059 erythrocyte genes with a text-mining score over 0.25, a threshold chosen because all six of our exemplar loci surpass it on both lists (Supp Tables 1, 2, and 3). We also examined higher thresholds to filter out weakly-associated genes and false positives (Figure 5). As a control, we tested for enrichment in four other cell types: hepatocytes, neurons, epithelial cells, and endothelial cells. To look for malaria-associated polymorphisms, we compared our top outliers against the top loci for malaria susceptibility in three independent GWAS, regardless of whether they were significant. The first study (Timmann et al. 2012) reports 50 variants with p < 5e-05; we considered all genes within 100kb of these variants, excluding genes adjacent to *HBB* and *ABO* as these adjacent genes are unlikely to be causal. The second study (Malaria Genomic Epidemiology Network, 2019) reports 97 genomic regions overlapping variants with a Bayes factor> 1000; we considered all genes within these regions, excluding genes in the regions overlapping *HBB*, *ABO*, *GYPA/B/E*, and HLA, other than the exemplar loci themselves, as the other genes in those regions are unlikely to be causal. The third study (Milet et al. 2019) reports 28 genes with p < 1e-05; we considered all of these genes. To look for transmembrane domains, we used TMHMM v. 2.0 (Sonnhammer et al. 1998). To look for *Plasmodium*- or Piroplasm-interacting proteins, we used the curated list of Ebel et al. (2017). To look for low-recombination genes, we used a genetic map (Hirsh et al. 2011) to calculate recombination rate in overlapping windows of 100-200 kb, and we defined low-recombination regions as those averaging less than 0.01 cM/Mb, which overlap 1054 genes.

## Acknowledgements

This project benefited from helpful insights provided by Lucas Buyon, Angela Early, Jonathan Goldberg, Usheer Kanjee, Dan Neafsey, Varshini Odayar, Alexa Rome, Estela Shabani, April Wei, and others. This work was supported by National Institutes of Health grants HL139337 and AI140751.

## Supplementary Tables

**Supp Table 1.**
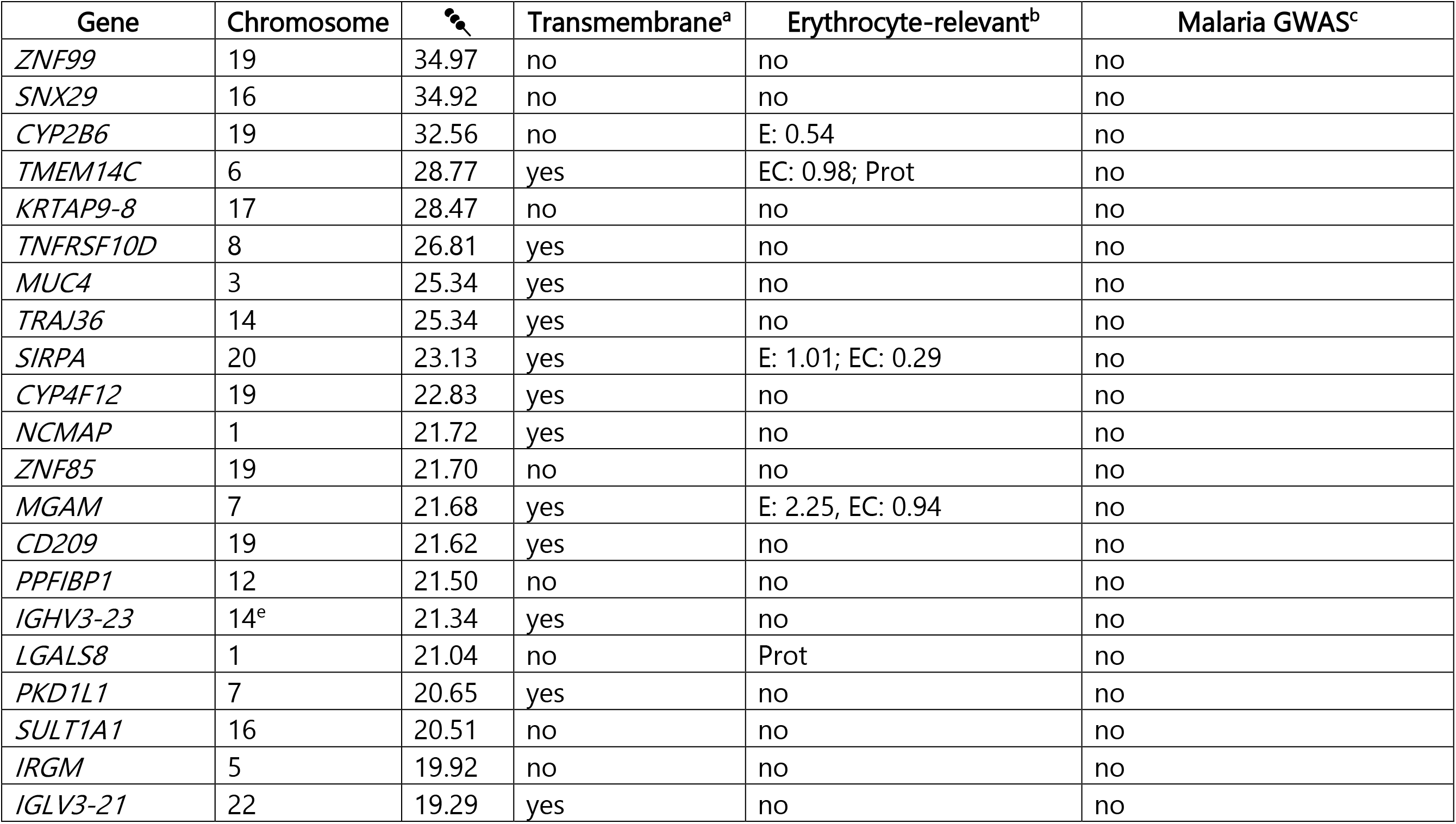
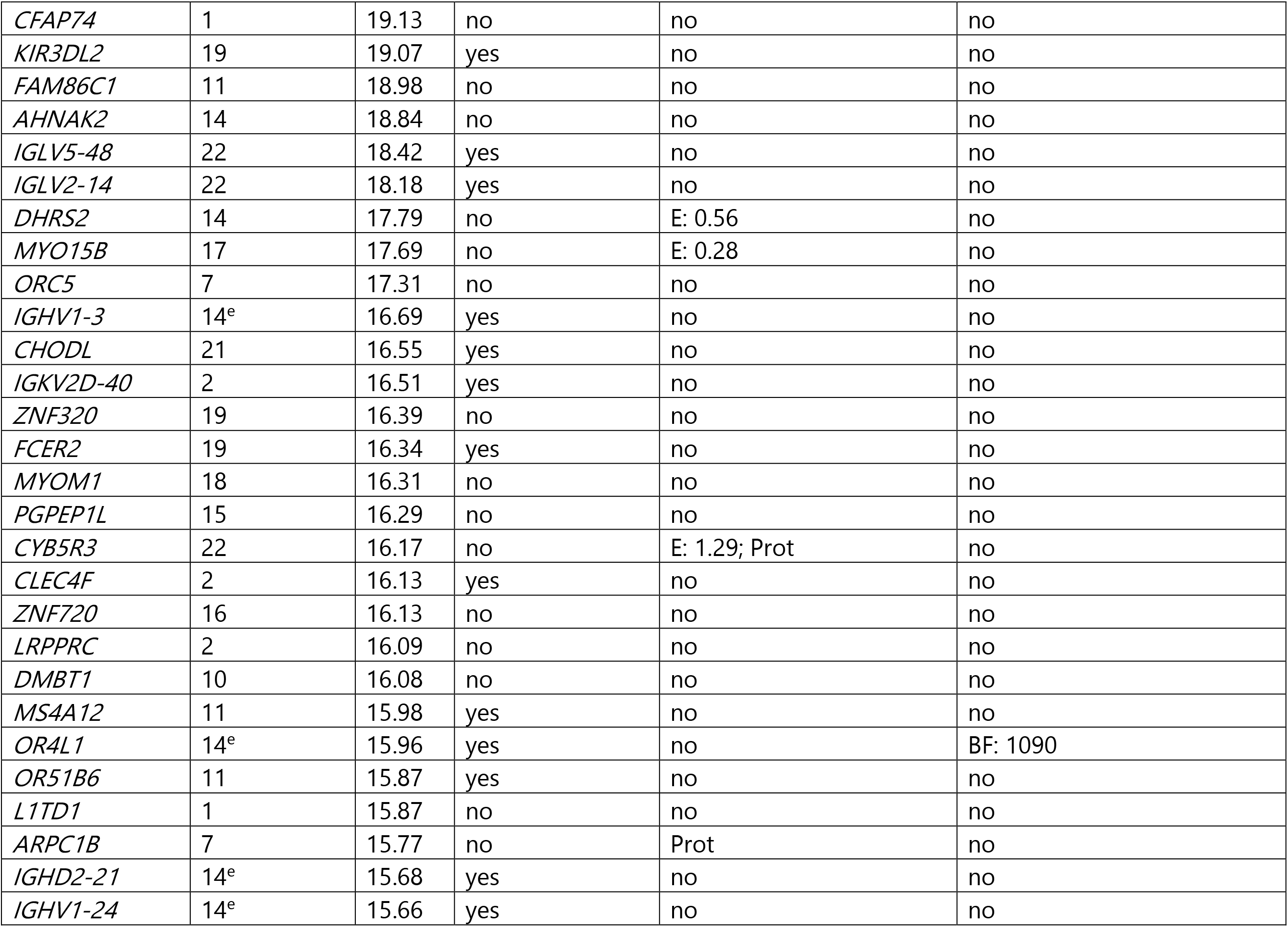
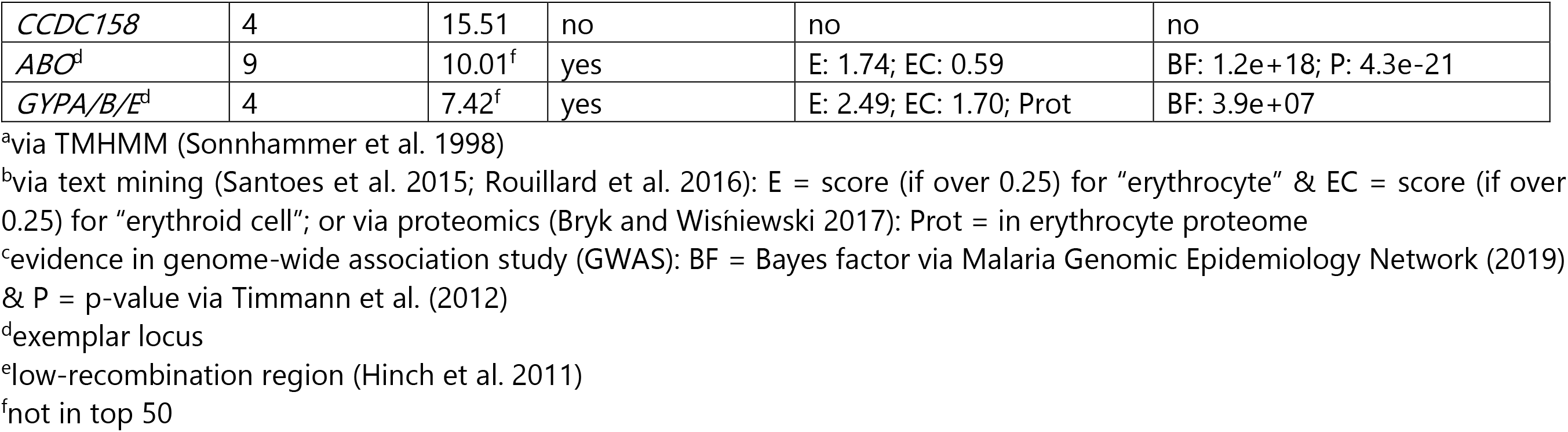
Top 50 genes based on 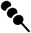 in exons or within 1kb upstream, excluding the HLA region, along with the two exemplar loci. *g* = 500 bp.

**Supp Table 2.** Top 50 genes based on 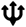 in exons or within 1kb upstream, along with the two exemplar loci. *g* =5000 bp.

| Gene | Chromosome | $\psi$ | Transmembrane<sup>a</sup> | Erythrocyte-relevant<sup>b</sup> | Malaria GWAS<sup>c</sup> |
| --- | --- | --- | --- | --- | --- |
| *PTK6* | 20 | 1.54e-08 | no | no | no |
| *SRMS* | 20 | 3.42e-07 | no | no | no |
| *FBXO31* | 16 | 8.85e-07 | no | no | no |
| *SPNS2* | 17 | 9.10e-07 | yes | E: 0.26 | BF: 3020 |
| *TTLL10* | 1<sup>e</sup> | 1.03e-06 | no | no | no |
| *IGHG1* | 14<sup>e</sup> | 7.02e-05 | yes | Prot | no |
| *SEMA4C* | 2 | 8.87e-06 | yes | no | no |
| *FAM178B* | 2 | 8.87e-06 | no | no | no |
| *MAP1LC3B* | 16 | 1.10e-05 | no | Kang et al. 2012 | no |
| *ZCCHC14* | 16 | 1.10e-05 | no | no | no |
| *OCA2* | 15 | 1.53e-05 | yes | no | no |
| *CCDC92B* | 17 | 1.78e-05 | no | no | no |
| *PPDPF* | 20 | 1.87e-05 | no | no | no |
| *PRKAG3* | 2 | 2.95e-05 | no | no | no |
| *GSG1L2* | 17 | 3.11e-05 | yes | no | no |
| *SORD* | 15 | 3.35e-05 | no | E: 1.17; Prot | no |
| *SLC35F3* | 1 | 3.50e-05 | yes | no | no |
| *GLP2R* | 17 | 4.54e-05 | yes | no | no |
| *TMEM88B* | 1<sup>e</sup> | 4.94e-05 | yes | no | no |
| *ANKRD65* | 1 | 4.94e-05 | no | no | no |
| *RSRP1* | 1 | 5.83e-05 | no | no | no |
| *RHD* | 1 | 5.83e-05 | yes | E: 2.18; EC: 0.85; Prot | no |
| *BRF1* | 14 | 7.23e-05 | no | no | no |
| *MLLT3* | 9 | 7.31e-05 | no | no | no |
| *EEFSEC* | 3 | 7.76e-05 | no | Prot | no |
| *CHRM5* | 15 | 9.22e-05 | yes | no | BF: 1980 |
| *SNX20* | 16 | 9.47e-05 | no | no | no |
| *GALNT18* | 11 | 1.02e-04 | yes | no | no |
| *CNTN2* | 1 | 1.21e-04 | no | no | no |
| *SPIN1* | 9 | 1.25e-04 | no | no | no |
| *CNNM4* | 2 | 1.35e-04 | yes | no | no |
| *HELZ2* | 20 | 1.41e-04 | no | no | no |
| *FNDC11* | 20 | 1.41e-04 | no | no | no |
| *METTL7B* | 12 | 1.41e-04 | yes | no | BF: 1030 |
| *ITGA7* | 12 | 1.41e-04 | yes | no | BF: 1030 |
| *HERC2* | 15 | 1.50e-04 | no | no | no |
| *ANKRD39* | 2 | 1.55e-04 | no | no | no |
| *ANKRD23* | 2 | 1.55e-04 | no | no | no |
| *ARHGAP26* | 5 | 1.60e-04 | no | no | no |
| *CD5* | 11 | 1.60e-04 | yes | E: 0.90 | no |
| *MYH9* | 22 | 1.69e-04 | no | Smith et al. 2019; Prot | no |
| *MOGS* | 2 | 1.73e-04 | yes | Prot | no |
| *MRPL53* | 2 | 1.73e-04 | no | no | no |
| *CCDC142* | 2 | 1.73e-04 | no | no | no |
| *PTPRM* | 18 | 1.73e-04 | yes | no | P: 3.8e-08 |
| *SFTA3* | 14 | 1.74e-04 | no | no | no |
| *ADCYAP1R1* | 7 | 1.93e-04 | yes | no | no |
| *MYLK4* | 6 | 1.94e-04 | no | no | P: 3.6e-07 |
| *TBC1D32* | 6 | 1.99e-04 | no | no | no |
| *MRPL20* | 1<sup>e</sup> | 2.00e-04 | no | no | no |
| *ACKR1*<sup>d</sup> | 1 | 4.35e-04<sup>f</sup> | yes | E: 1.99; EC: 0.34; Prot | not in the included studies |
| *CR1*<sup>d</sup> | 1 | 5.65e-03<sup>f</sup> | yes | E: 2.50; EC: 0.40; Prot | not in the included studies |

<sup>a</sup>via TMHMM (Sonnhammer et al. 1998)
<sup>b</sup>via text mining (Santoès et al. 2015; Rouillard et al. 2016): E = score (if over 0.25) for “erythrocyte” & EC = score (if over 0.25) for “erythroid cell”; or a reference supporting erythroid relevance even without a high text mining score; or via proteomics (Bryk and Wisniewski 2017): Prot = in erythrocyte proteome
<sup>c</sup>evidence in genome-wide association study (GWAS): BF = Bayes factor via Malaria Genomic Epidemiology Network (2019) & P = p-value via Milet et al. (2019)
<sup>d</sup>exemplar locus
<sup>e</sup>low-recombination region (Hinch et al. 2011)
<sup>f</sup>not in top 50

**Supp Table 3.** Top 50 genes based on 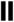 in exons or within 1kb upstream, excluding the HLA region, along with the two exemplar loci.

| Gene | Chromosome | $\Pi$ | Transmembrane<sup>a</sup> | Erythrocyte-relevant<sup>b</sup> | Malaria GWAS<sup>c</sup> |
| --- | --- | --- | --- | --- | --- |
| *RLIM* | X | 495.12 | no | no | no |
| *ABCB7* | X<sup>e</sup> | 272.91 | yes | E: 0.26; EC: 1.42 | no |
| *HBB*<sup>d</sup> | 11 | 194.83 | no | E: 2.46; EC: 2.06; Prot | BF: 1.8e+69; P: 5.6e-14 |
| *UGT2B10* | 4 | 187.49 | yes | no | no |
| *RABGAP1L* | 1<sup>e</sup> | 162.56 | no | EC: 0.86; Prot | no |
| *ZDHHC15* | X<sup>e</sup> | 162.45 | yes | no | no |
| *ZKSCAN4* | 6 | 162.19 | no | no | no |
| *ZSCAN16* | 6<sup>e</sup> | 153.98 | no | EC: 0.80 | no |
| *ZKSCAN8* | 6 | 153.98 | no | no | no |
| *GABBR1* | 6 | 152.81 | yes | no | no |
| *OR5V1* | 6 | 143.49 | yes | no | no |
| *TARDBP* | 1 | 141.49 | no | Prot | no |
| *RABGEF1* | 7<sup>e</sup> | 141.35 | no | Prot | no |
| *CSMD3* | 8 | 139.00 | yes | no | no |
| *TESK2* | 1 | 136.32 | no | no | no |
| *EIF2B3* | 1 | 135.39 | no | Prot | no |
| *PRDX1* | 1 | 135.37 | no | E: 0.35; Prot | no |
| *UROD* | 1 | 130.54 | no | E: 1.79; Prot | no |
| *HECTD3* | 1 | 130.54 | no | Prot | no |
| *MAS1L* | 6 | 129.21 | yes | no | no |
| *OR2B3* | 6<sup>e</sup> | 129.07 | yes | no | no |
| *ZSWIM5* | 1 | 128.42 | no | no | no |
| *ZSCAN12* | 6<sup>e</sup> | 125.75 | no | no | no |
| *KRTAP4-4* | 17 | 125.66 | no | no | no |
| *DNAH14* | 1<sup>e</sup> | 123.89 | no | Prot | no |
| *OIT3* | 10 | 122.99 | no | no | no |
| *NKAPL* | 6<sup>e</sup> | 122.91 | no | no | no |
| *UGT2B4* | 4 | 122.69 | yes | no | no |
| *SLC9C2* | 1 | 122.20 | yes | no | no |
| *ABCC12* | 16 | 121.65 | yes | no | no |
| *SPAG16* | 2 | 121.48 | no | no | no |
| *ZSCAN23* | 6 | 121.30 | no | no | no |
| *ZBTB37* | 1 | 119.20 | no | no | no |
| *SLC38A4* | 12 | 118.48 | yes | no | no |
| *ANKRD45* | 1 | 116.10 | no | no | no |
| *KRT32* | 17 | 114.66 | no | Prot | no |
| *MOG* | 6 | 113.55 | no | no | no |
| *CNTLN* | 9 | 113.05 | no | no | no |
| *QRICH1* | 3 | 112.89 | no | no | no |
| *VKORC1L1* | 7 | 111.13 | yes | Prot | no |
| *PLA2G12B* | 10 | 106.82 | no | no | no |
| *KRTAP2-2* | 17 | 106.65 | no | no | no |
| *NIPBL* | 5<sup>e</sup> | 105.43 | no | no | BF: 3170 |
| *CPLANE1* | 5 | 105.33 | no | no | no |
| *MBNL3* | X | 103.46 | no | no | no |
| *GPX2* | 14 | 103.42 | no | no | no |
| *RC3H1* | 1<sup>e</sup> | 102.85 | no | no | no |
| *KRT35* | 17 | 101.02 | no | Prot | no |
| *KDM3A* | 2 | 100.31 | no | Prot | no |
| *KRTAP2-3* | 17 | 99.97 | no | no | no |
| *G6PD*<sup>d</sup> | X | 68.80<sup>f</sup> | no | E: 2.97; EC: 0.58; Prot | not in the included studies |

<sup>a</sup>via TMHMM (Sonnhammer et al. 1998)
<sup>b</sup>via text mining (Santoès et al. 2015; Rouillard et al. 2016): E = score (if over 0.25) for "erythrocyte" & EC = score (if over 0.25) for "erythroid cell"; or via proteomics (Bryk and Wisniewski 2017): Prot = in erythrocyte proteome
<sup>c</sup>evidence in genome-wide association study (GWAS): BF = Bayes factor via Malaria Genomic Epidemiology Network (2019) & P = p-value via Timmann et al. (2012)
<sup>d</sup>exemplar locus
<sup>e</sup>low-recombination region (Hinch et al. 2011)
<sup>f</sup>not in top 50

**Supp Table 4.**
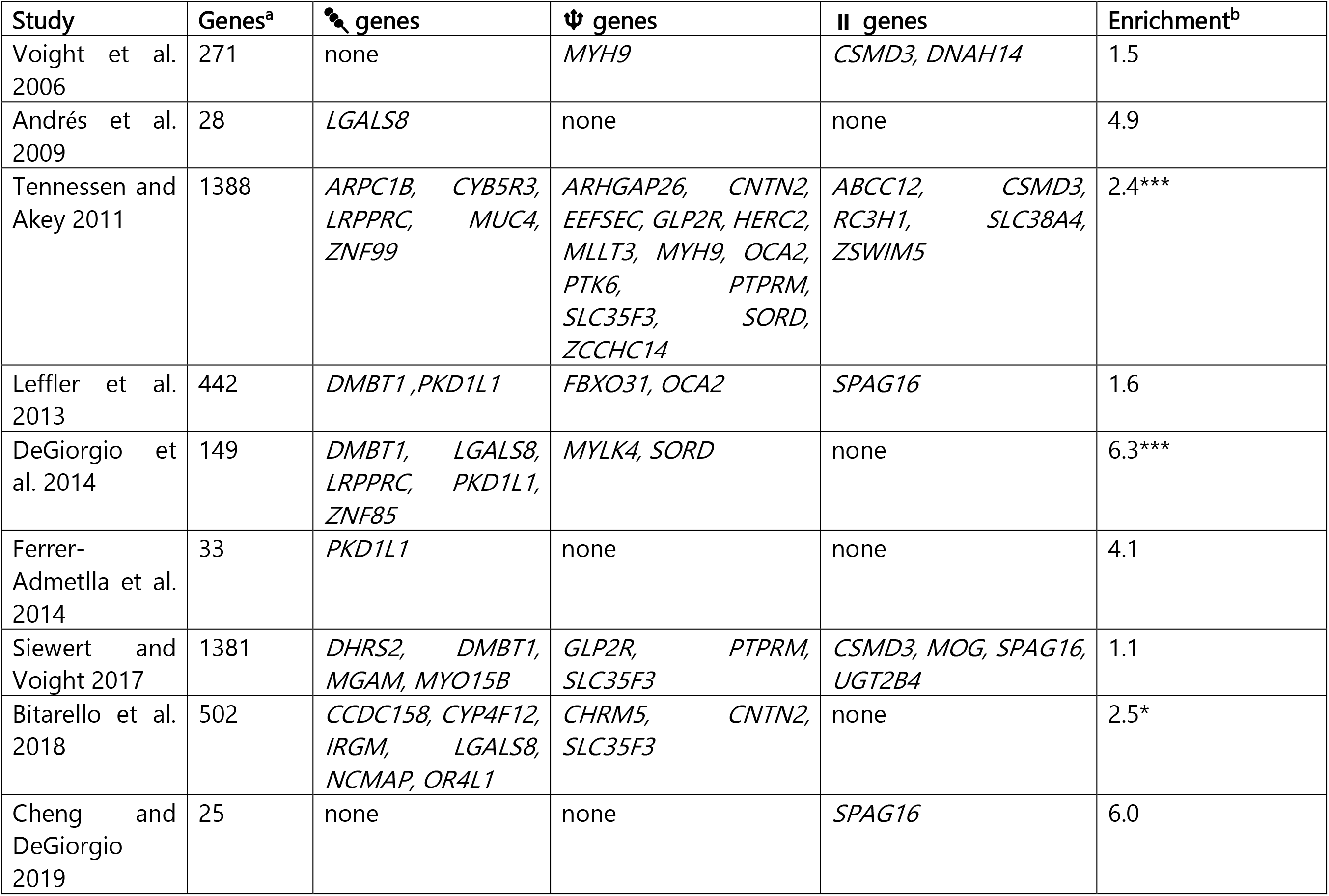
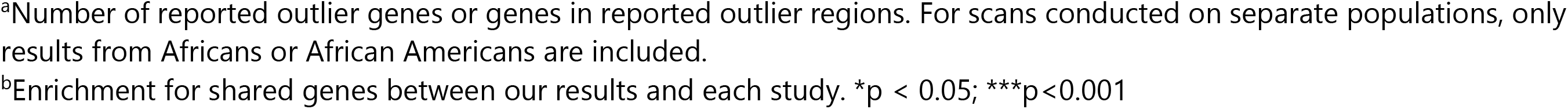
Overlap between outliers in this study and those from other genome-wide scans for selection.

## Supplementary Figures

**Supplementary Figure 1.**
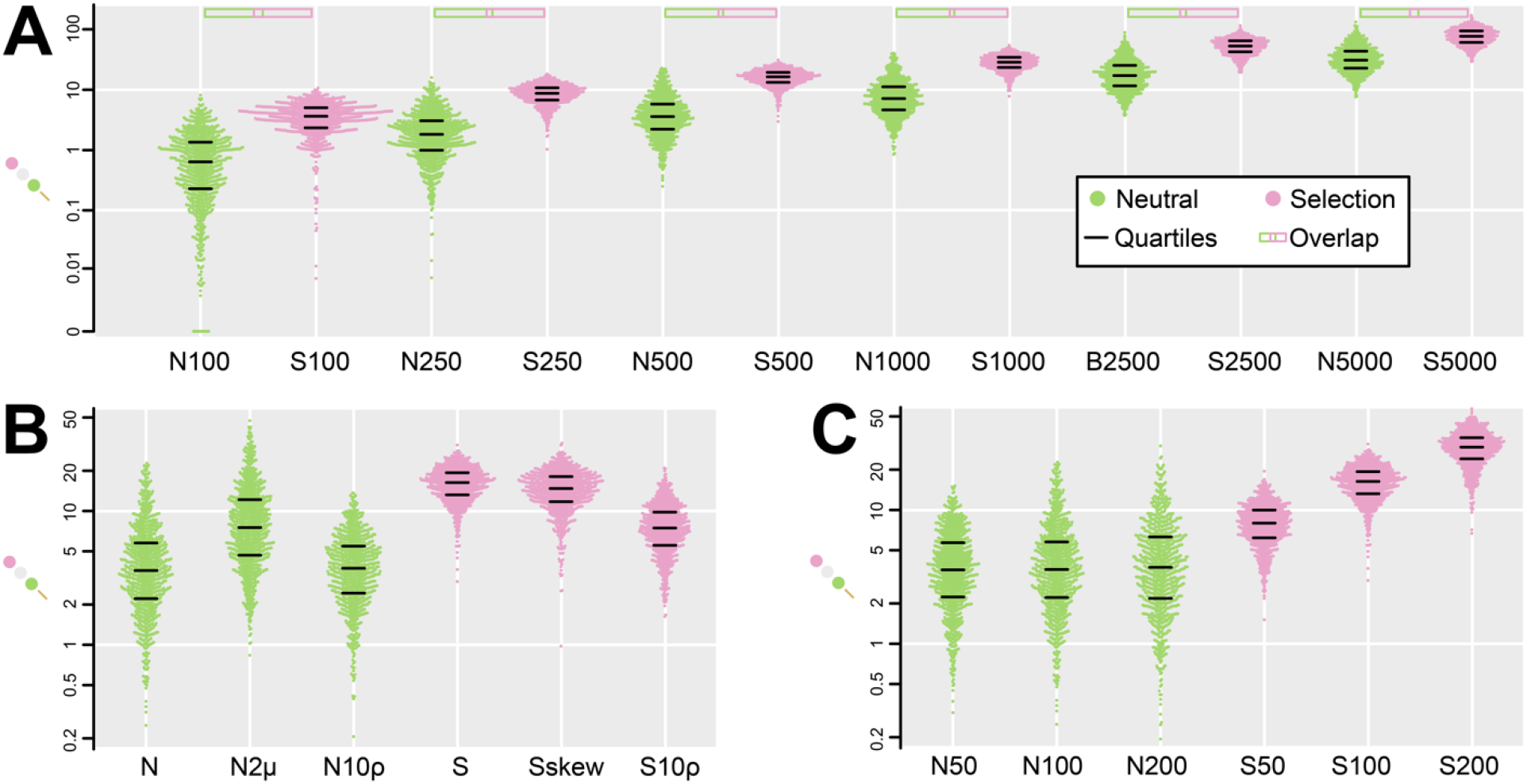
Simulation results for 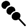. (A). Distributions of 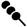 values for neutral and selection scenarios with varying *g* values, with ρ = 0.001, expected MAF = 0.5, μ = 1e-7, and 100,000 generations. Distributions are labeled N (“neutral”) or S (“selection”), along with *g* value (e.g. N250 = neutral simulations with *g* = 250 bp). Boxes above each pair of neutral and selection distributions indicate the degree of overlap between the distributions. Overlap is minimized for intermediate *g* values of 500 and 1000 bp (lower 7% S quantile meets upper 7% N quantile). (B) Distributions of 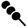 values for neutral and selection scenarios with *g* = 500 bp and 100,000 generations. “N” and “S” are the same as “N500” and “S500” in (A), and the remaining distributions have the same parameters except for a single alteration each. “N2μ” has a doubled mutation rate, “N10ρ”and “S10ρ” have 10-fold increased recombination rate, and “Sskew” has expected MAF of 0.08. (C). Distributions of 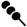 values with *g* = 500bp, ρ = 0.001, expected MAF = 0.5, μ = 1e-7, and age of the balanced polymorphism either 50,000 (N50 and S50), 100,000 (N100 and S100) or 200,000 (N200 and S200) generations.

**Supplementary Figure 2.**
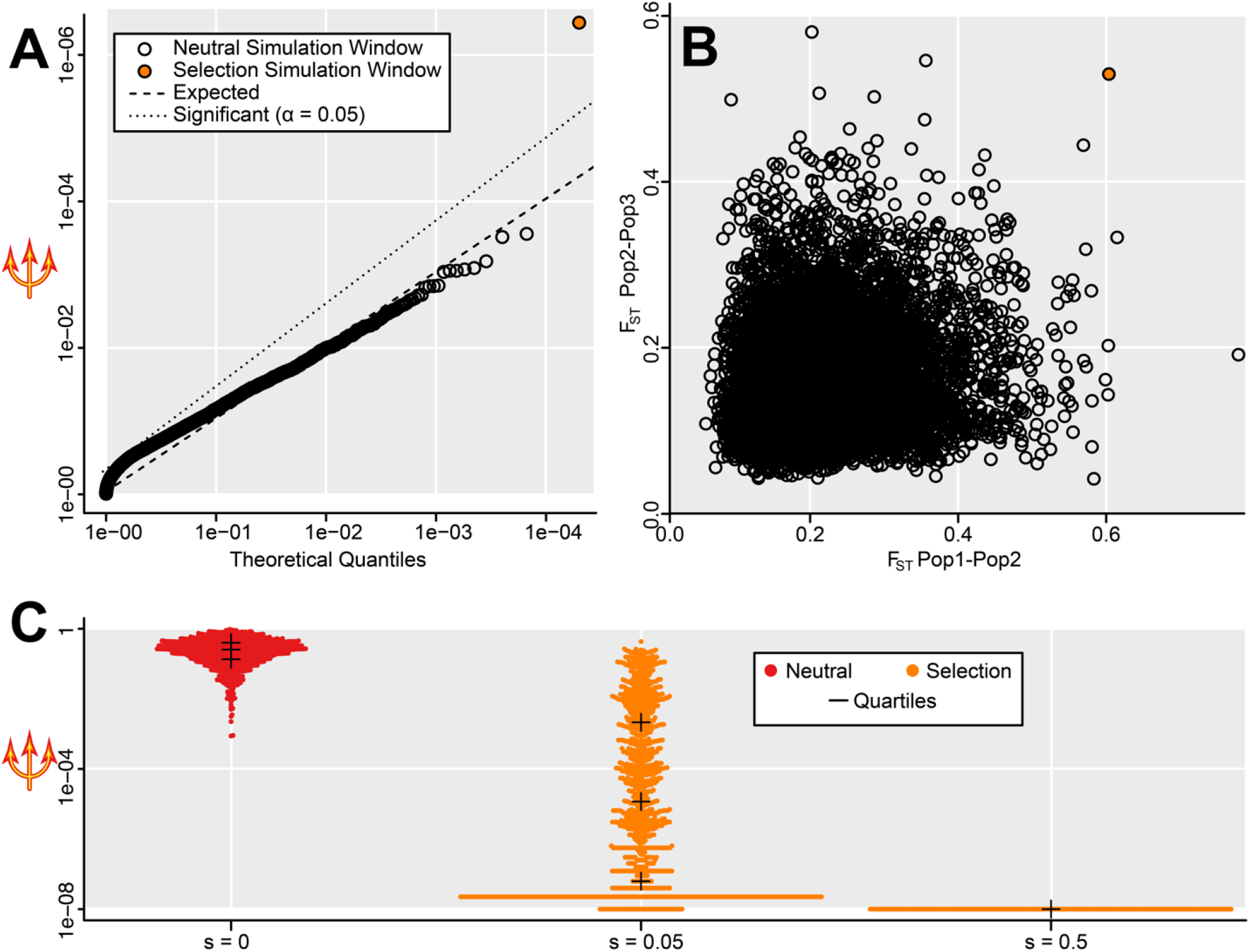
Simulation results for 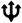. (A) Out of 10,000 neutral windows and a single selection window, the selection window is the clear significant outlier in a QQ plot. (B) In this case, the selection outlier does not show the highest F_ST_ in any comparison, yet it is an extreme outlier for 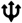 which combines across F_ST_ values. (C) Distribution of 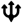 across 1000 windows each with *s* of 0, 0.05, or 0.5, when each window is included alone among 10,000 neutral windows. 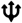 is substantially lower when selection operates. Every window with *s* = 0.5 shows the lowest possible value of 1e-08, so these points are depicted as overlapping rather than fully spread out.

**Supplementary Figure 3.**
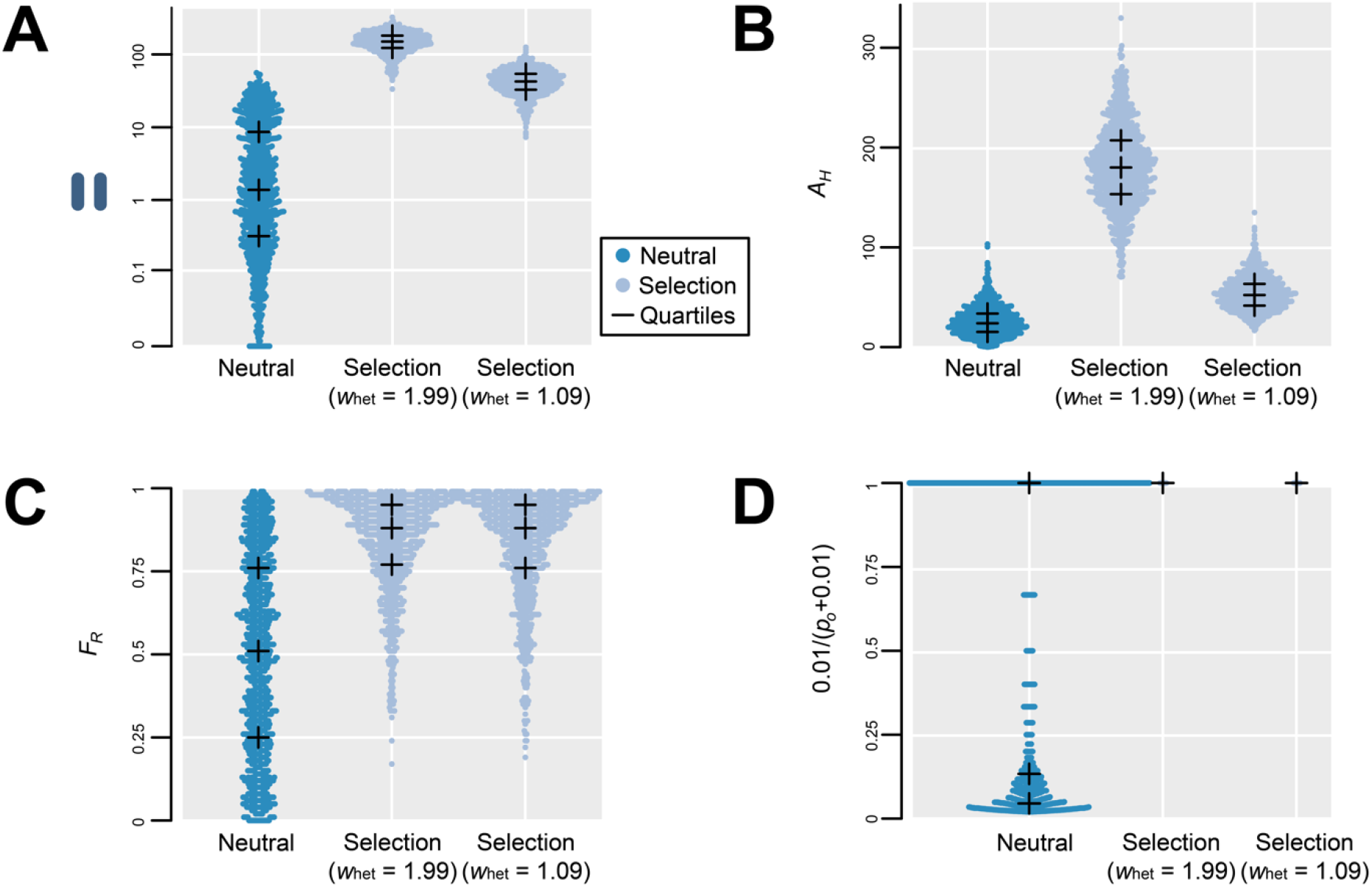
Simulation results for 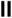. (A) Distributions of 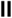 values for three evolutionary scenarios: neutral evolution, strong selection (heterozygote fitness *w_het_* = 1.99), and weak selection (heterozygote fitness *w_het_* = 1.09). (B) Distributions of the difference in total heterozygous sites times ingroup allele frequency (*A_H_*) for the same evolutionary scenarios. (C) Distributions of F_ST_ rank proportion (*F_R_*) for the same evolutionary scenarios. (D) Distributions of the adjusted reciprocal of the outgroup allele frequency (0.01/(*p_o_*+0.01) for the same evolutionary scenarios; selection scenarios are always at one because *p_o_* is zero, so datapoints are overlapped for ease of visualization.

